# Benchmarking virus identification tools on vaginal metagenomes

**DOI:** 10.64898/2026.09.24.753817

**Authors:** Faruk Dube, Anna-Ursula Happel, Heather B. Jaspan, Luisa Warchavchik Hugerth

## Abstract

The vaginal microbiome is associated with reproductive and sexual health, yet its viral component remains understudied. A recent vaginal genome catalogue reported that 85.8% of its species-level viral groups were absent from all five viral databases examined. Viral identification tools depend on reference sequences, either directly or through their training data. Previous benchmarks used simulated or non-vaginal datasets. How well these tools generalise to vaginal metagenomes therefore remains unclear. We benchmarked 14 tools spanning five methodological approaches. Evaluations used controlled genome fragments, viral spike-in co-assemblies and virus-like particle (VLP)-enriched co-assemblies. A multi-evidence benchmark assessed 13 vaginal shotgun assemblies (primary cohort). An independent 30-sample cohort provided external validation. geNomad ranked first or second by Matthews correlation coefficient (MCC) in every ranked evaluation. VIBRANT achieved similar MCC to geNomad on the primary assemblies. It made eight false-positive calls among 11,577 negatives but recovered only 22% of viral contigs. VIBRANT ranked first in the independent cohort. Tool rankings agreed closely between cohorts (Spearman ρ = 0.952; 95% CI 0.921–0.960). Detection of sequences lacking reference homologues differed substantially between tools. Two phage genomes without detectable reference homologues provided a controlled novelty test. Each evaluated composition-based tool recovered at least 74% of their 1,500-bp fragments. Marker-or reference-dependent tools recovered at most 31%. A separate analysis examined 48 unconfirmed candidate viral contigs lacking detectable nucleotide or protein homologues. Jaeger flagged 41, compared with four for VIBRANT and one for geNomad. On short fragments, attention-based tools achieved 4.0–5.5 times the mean area under the precision–recall curve of feature-based tools. ViraLM accounted for most of this difference. ViraLM also had the highest recall (sensitivity) of eukaryote-infecting viruses in the evaluated panel. VirSorter2 achieved the highest MCC on VLP-enriched contigs. These results support geNomad when prioritising overall classification performance and VIBRANT when minimising false-positive phage calls is the priority. ViraLM supports short-contig and eukaryote-infecting-virus detection. VirSorter2 supports identification in VLP-enriched assemblies, and Jaeger supports screening for candidate novel phages. These findings guide tool selection for studies investigating the vaginal virome in reproductive and sexual health.

## Introduction

The composition of the vaginal microbiome (VMB) is associated with reproductive and sexual health outcomes, including bacterial vaginosis (BV), preterm birth, and susceptibility to sexually transmitted infections [1–4]. Research has focused primarily on its bacterial component, leaving the virome comparatively understudied [5–7]. The virome is composed predominantly of bacteriophages (hereafter phages) and eukaryote-infecting viruses [8, 9]. Phages have been associated with BV status [8, 9] and with vaginal community dynamics [10]. Eukaryote-infecting viruses, while less abundant, include clinically important pathogens and also interact with the bacterial community [7]. A systematic review linked vaginal bacterial community composition with human papillomavirus (HPV) infection and cervical dysplasia [11]. Investigating these relationships requires reliable identification of viral sequences in vaginal metagenomes.

Reference coverage is a concern for vaginal virus identification due to the underrepresentation of vaginal viral sequences in major databases. The Integrated Microbial Genomes Viral database (IMG/VR) and its successor MetaVR are the most comprehensive metagenomic viral references available [12, 13]. In MetaVR, the vaginal category accounts for 0.27% of uncultivated viral genomes (UViGs) assigned to human-associated ecosystems (MetaVR ecosystem statistics, accessed 2026-09-14). This gap is further illustrated in the Vaginal Microbial Genome Collection (VMGC) [14]. In this catalogue, 85.8% of species-level viral groups were absent from all five viral databases examined. Resources such as VMGC and VIRGO2 are expanding the available vaginal genome and gene catalogues [15]. However, database coverage alone does not establish whether identification tools can detect these sequences. Species-level absence also does not establish what each tool encountered during training. Direct evaluation is therefore needed to assess detection of vaginal viruses with limited reference similarity.

We group these tools into five methodological approaches: reference-based methods (MetaPhinder and Sourmash; [16, 17]), feature-based machine learning (VirSorter, VirSorter2, VIBRANT, and VirFinder; [18–21]), convolutional and recurrent neural networks (DeepVirFinder, PPR-Meta, Jaeger, HVSeeker, and Seeker; [22–26]), attention-based transformer architectures (ViraLM and TransGINmer; [27, 28]), and hybrid deep-learning-plus-marker approaches (geNomad; [29]).

Several biological contexts warrant consideration when comparing these tools. Prophage-like elements have been recovered from cervicovaginal metagenomes of South African adolescents [30]. These findings motivate screening bacterial contigs for prophage regions and distinguishing viral from flanking bacterial sequence [31]. Vaginal bacterial communities also differ across community state types (CSTs). CSTs I, II, III, and V are dominated by individual *Lactobacillus* species, whereas CST-IV encompasses more diverse communities containing BV-associated bacteria [32, 33]. Bacterial community composition has been associated with phage profiles [8, 34]. These associations motivate examining identification performance across community backgrounds.

Previous benchmarks, built on simulated datasets and non-vaginal metagenomes, have shown that tool performance varies [35–38]. Other studies also examined agreement between tools and strategies for combining their predictions [39, 40]. These evaluations do not establish how the tools perform in vaginal metagenomes. Feature-based tools score annotated genes, and short contigs carry few genes. Attention-based models learn sequence context directly and may retain accuracy on short contigs. Tools also differ in their declared target scope, and many were designed mainly for phages. Whether declared scope predicts detection of eukaryote-infecting viruses has not been tested. We therefore addressed four questions. Do attention-based tools outperform feature-based machine learning tools on short contigs (Q1)? Does a tool’s declared target scope predict detection of eukaryote-infecting viruses (Q2)? How does detection change with sequence novelty relative to reference databases (Q3)? Do marker-dependent tools have an advantage on prophage-containing contigs (Q4)?

Here, we benchmarked 14 viral identification tools spanning five methodological approaches on vaginal metagenomes. We evaluated the tools on three benchmark tracks and a multi-evidence benchmark. Track A addressed Q1–Q3 using controlled fragments of viral and bacterial genomes at six lengths. Track B used viral spike-in co-assemblies across coverage levels and community backgrounds. Track C used co-assemblies of matched shotgun and virus-like particle (VLP)-enriched reads. The multi-evidence benchmark evaluated 13 vaginal shotgun assemblies and included prophage-containing contigs to address Q4. An independent 30-sample cohort provided external validation. Beyond Q1–Q4, we assessed performance across assembly coverage and community backgrounds. We also examined the consistency of tool rankings across evaluations and cohorts. A separate analysis compared predictions for candidate viral contigs lacking detectable nucleotide or protein homologues. Their viral identity remained unconfirmed. We also compared computational costs. Our aim was to guide tool selection by clarifying trade-offs between viral recovery and false-positive calls in vaginal metagenomes.

## Results

The three tracks and the multi-evidence benchmark provided four complementary evaluation sets (**Figure 1**). Track A included 421 viral and 6,268 bacterial fragments at 1,500 bp (1:15; **Table S1**), with corresponding panels at six lengths (500 bp–10,000 bp). Track B combined a six-depth coverage sweep in two backgrounds with a 10× diversity contrast across four CST-I and four CST-IV-B backgrounds. Track C contained 29 viral positives and 931 non-viral negatives defined by enrichment and annotation (1:32; **Table S35**), from 13 matched shotgun–virome sample pairs representing 11 participants. Additionally, the multi-evidence benchmark retained 1,453 viral positives and 11,577 grounded negatives from 13 shotgun assemblies (13,030 contigs ≥ 1,500 bp; **Figure S14**). The independent cohort contributed 30 shotgun metagenomes.

**Figure 1.**
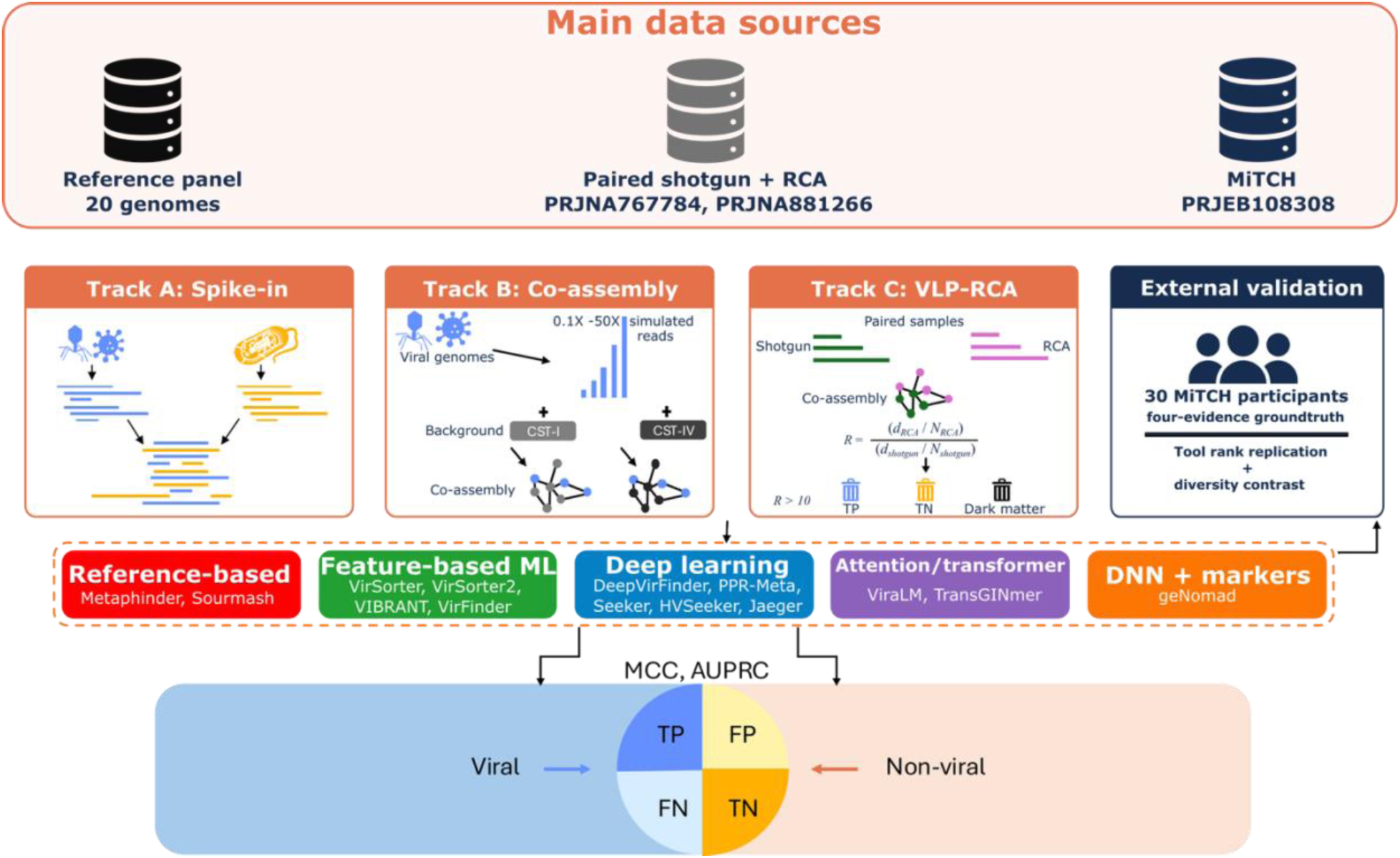
A reference genome panel and two vaginal cohorts feed three benchmark tracks and an independent validation of viral identification tools. Three data sources supply the complementary benchmarks: a 20-genome viral and bacterial reference panel, the paired shotgun-plus-RCA UChoose cohort (13 samples; PRJNA767784, PRJNA881266), and the 30-sample MiTCH cohort (PRJEB108308). Track A fragments the reference genomes into controlled spike-ins; Track B co-assembles simulated viral reads (0.1×–50×) with real CST-I and CST-IV-B vaginal metagenome backgrounds; and Track C co-assembles matched UChoose shotgun and VLP-RCA reads, labelling contigs by the library-size-normalised phi29 enrichment ratio *R* (true positive, true negative, dark matter). The same 14 tools, grouped into reference-based, feature-based machine-learning, deep-learning, attention/transformer, and Deep Neural Network-plus-marker approaches, are scored by MCC and AUPRC from viral/non-viral confusion matrices. External validation applies a four-evidence ground truth to the 30 MiTCH shotgun metagenomes to test tool-rank replication and a community-diversity contrast.

The Matthews correlation coefficient (MCC) ranges from −1 (complete disagreement) to 1 (perfect classification) [41]. The area under the precision–recall curve (AUPRC) summarises performance across score thresholds.

### geNomad achieves the highest classification accuracy on controlled fragments

On Track A at 1,500 bp, geNomad achieved the highest MCC of 0.651 [95% CI 0.613 to 0.690]. ViraLM was the only other tool above MCC 0.3 (0.536 [95% CI 0.508 to 0.563]), with VirSorter2 third at 0.283 [95% CI 0.237 to 0.326] (**Figure 2A**; **Table S4**). The top two tools reversed on AUPRC, with ViraLM at 0.811 [95% CI 0.776 to 0.841] and geNomad at 0.484 [95% CI 0.438 to 0.532]. These leaders occupied opposite operating points, with geNomad favoring precision and ViraLM recall (**Figure 2B**; **Figure S2** for the same comparison at 3,000 bp; **Table S34**). The false-discovery rate was 0.645 for ViraLM and 0.103 for geNomad (**Table S34**). In-sample threshold optimisation increased ViraLM’s MCC from 0.536 to 0.698, TransGINmer’s from 0.280 to 0.355, and PPR-Meta’s from 0.149 to 0.279 (**Figure S4**; **Table S41**). These optimised scores were estimated on the same fragments used to select the thresholds.

**Figure 2.**
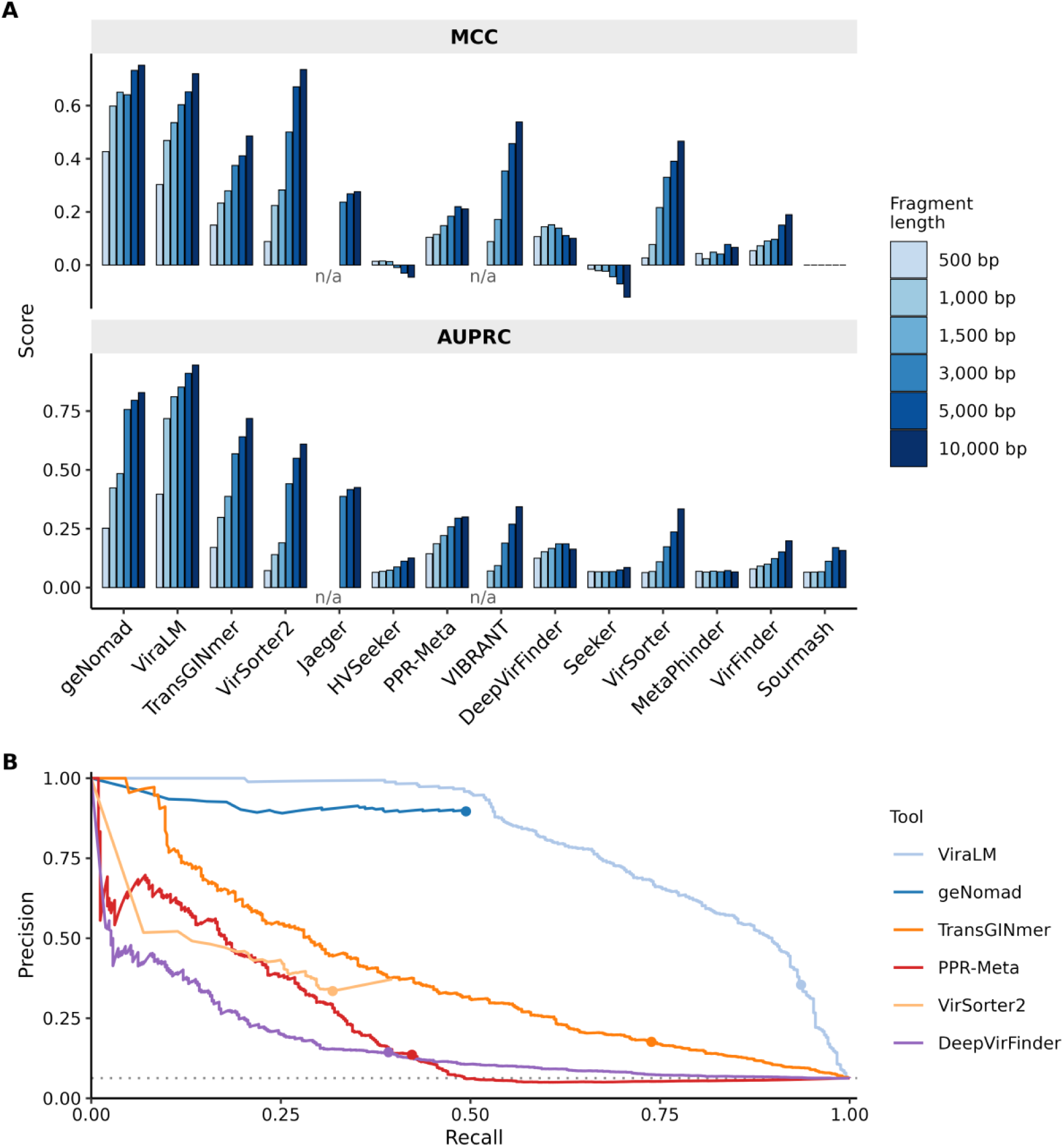
geNomad leads on Matthews correlation coefficient (MCC) while ViraLM leads on threshold-independent Area Under the Precision recall curve (AUPRC) at 1,500 bp. (**A**) MCC and AUPRC per tool across six Track A fragment lengths (500, 1,000, 1,500, 3,000, 5,000, 10,000 bp); 1,000-iteration bootstrap 95% confidence intervals for these estimates are given in **Table S4**. (**B**) Precision–recall curves for the six highest-AUPRC tools at 1,500 bp. Solid curves end at the last available score cutoff, without extrapolation. Filled circles mark the benchmark decision rules (**Table S21**), including p-value filtering for DeepVirFinder. The grey dotted line marks the random-classifier baseline at 6.3% viral prevalence. n = 421 viral and 6,268 bacterial fragments at 1,500 bp.

### Attention-based tools outperform feature-based ML tools on short fragments

On 500- and 1,000-bp fragments, each attention-based tool (ViraLM, TransGINmer) had higher AUPRC and recall than each evaluable feature-based ML tool (**Figure 3**; **Tables S4, S7**). Mean AUPRC was 4.0-fold higher at 500 bp (0.284 versus 0.072; VIBRANT produced no output) and 5.5-fold higher at 1,000 bp (0.508 versus 0.093). ViraLM contributed most of the difference (0.397 and 0.718). At 500 bp, feature-based AUPRC (0.064–0.079) was close to the 6.3% viral prevalence. Higher recall came with lower precision (ViraLM 0.19, TransGINmer 0.11 at 500 bp). geNomad, outside this comparison, had the highest MCC at both lengths (0.427 and 0.598). MCC rose with length for all six compared tools. The largest gains were for VirSorter2 (0.088 to 0.736 from 500 to 10,000 bp) and VIBRANT (0 to 0.538), and the smallest for the k-mer-based VirFinder (0.054 to 0.190). Among the remaining tools, HVSeeker’s MCC did not differ from zero at any length (−0.046 to 0.016). Seeker’s MCC was negative at every length, reaching −0.121 at 10,000 bp. Sourmash made no positive call (**Tables S4, S7**). Per-fragment correctness differed among tools at every fragment length (Cochran’s Q = 3,935 to 53,572, df = 13, all p < 0.001; **Table S5**). At 1,500 bp, 88 of 91 pairwise comparisons were significant after correction for multiple testing (McNemar’s tests, Benjamini–Hochberg q < 0.05; **Figure S3**; **Table S6**). Because 93.7% of fragments are bacterial, these correctness tests mainly reflect specificity, and Jaeger, which produces no output below 2,048 bp, was scored as calling every fragment negative; the three non-significant pairs were Jaeger versus Sourmash, neither of which made a positive call, and VIBRANT, which made 37 positive calls (**Table S34**), versus each of them. **Table S20** reports recall for each of the 14 viral genomes and false-positive calls for each of the 42 bacterial sequences. Those bacterial sequences are mostly draft-assembly contigs making up the six negative-control genomes.

**Figure 3.**
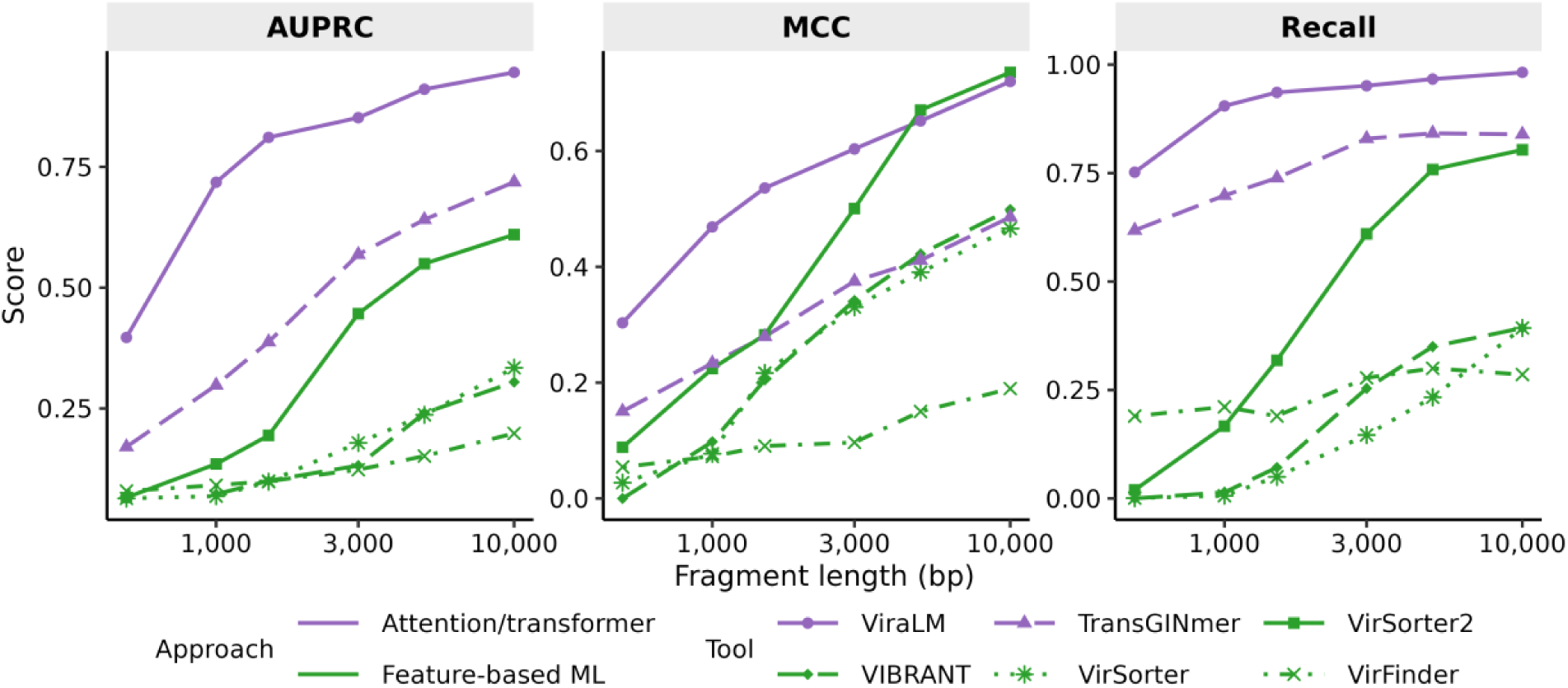
Attention-based architectures have superior performance for short fragments. AUPRC, MCC and recall versus fragment length for six attention-based or feature-based tools. Colour identifies approach; line type and point shape identify tools. Fragment length is log-spaced and y-axis ranges differ between metrics. Sample sizes: 6 tools across six fragment lengths from 500 to 10,000 bp; per-length fragment counts are in **Table S7**.

### Published scope alone is insufficient to guide eukaryote-infecting virus detection

Declared target scope did not predict eukaryote-infecting virus recall, which differed sharply between virus categories (**Figure 4**; **Figure S17**; **Tables S1, S21**). At 1,500 bp most tools recovered HPV fragments (mean recall 0.54, six tools ≥ 90%), whereas few recovered herpesvirus fragments (mean 0.26). Of the 204 herpesvirus fragments, from two herpes simplex virus genomes, recall exceeded 50% only for ViraLM (100%), TransGINmer (81%) and geNomad (60%). No other tool exceeded 27% (**Figure S5**). The fourth all-virus tool, VirSorter2, recovered 13% of herpesvirus and 25% of HPV fragments. Both figures are lower than those of the phage-only Seeker (27% and 100%) and HVSeeker (25% and 90%). At 3,000 bp, all four all-virus tools exceeded every other tool on herpesvirus recall (41–100% versus 0–22%). At both lengths, the tools declaring all-virus scope were also among the four highest-MCC tools. HPV recall did not track declared scope at either length. At 3,000 bp, Seeker, HVSeeker and PPR-Meta recovered all eight HPV fragments, as did the four all-virus tools. The phage-only Jaeger recovered seven at a specificity of 0.866. Seeker and HVSeeker had the panel’s two lowest specificities at 1,500 bp (0.442 and 0.483). ViraLM reached 100% recall in all three eukaryote-infecting categories at a specificity of 0.886 (**Table S34**). *Anellovirus* recall rests on two fragments from a single genome at 1,500 bp and one at 3,000 bp, too few to compare tools. Within the phage categories, recall also varied by host genus. At 3,000 bp Jaeger recovered 87.5–100% across the five genera, whereas HVSeeker ranged from 51.7% to 100% (**Figure 4**).

**Figure 4.**
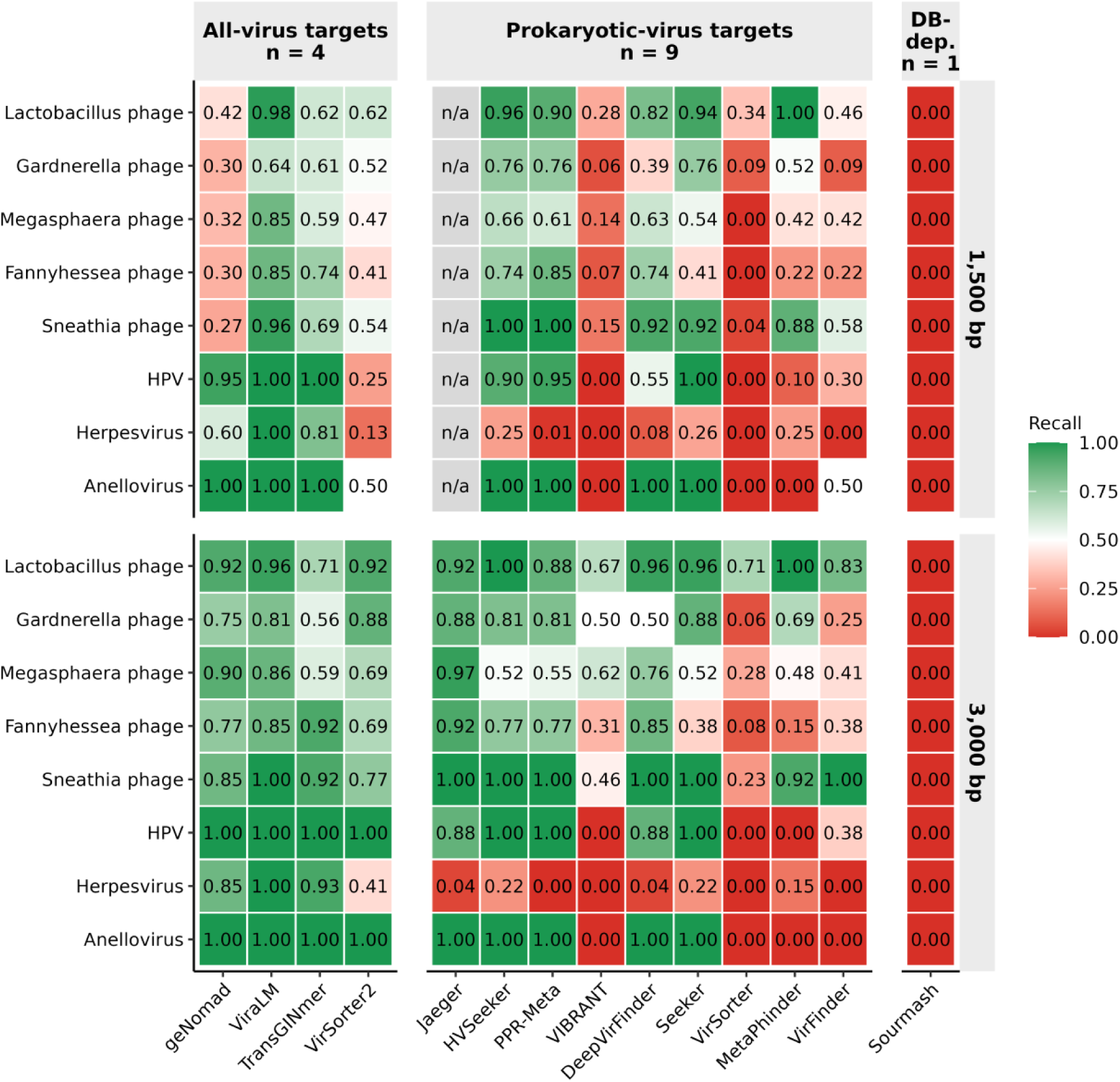
Declared target scope does not predict eukaryote-infecting virus recall. Recall by virus category for all 14 tools at 1,500 and 3,000 bp; phage categories identify bacterial host genera. Cell values and colour show recall from 0 (red) to 1 (green). Tools are grouped by declared target scope (**Table S21**); the prokaryotic-virus block pools phage-only, prokaryotic-virus and phage+plasmid tools, and DB-dep. is Sourmash, whose scope follows the user-supplied signature database. Grey n/a cells indicate Jaeger’s minimum input length exceeds 1,500 bp; 3,000 bp is the shortest Track A length at which all 14 tools are defined. Sample sizes: 421 viral fragments at 1,500 bp (2 anellovirus to 204 herpesvirus) and 205 at 3,000 bp (1 to 101), across eight categories.

### Recall on novel vaginal phage genomes diverges by homology dependence

Homology-free fragments distinguished tools by their dependence on external sequence resources. To address Q3, 15 vaginal phage genomes with < 95% average nucleotide identity (ANI) to MetaVR database were divided into 13 genomes with detectable relatives (84.3–94.9% BLASTn identity) and two with no detectable nucleotide or protein homology under our searches (**Figure 5**). At 1,500 bp, ViraLM and HVSeeker both recovered all 65 fragments from the two homology-free genomes. ViraLM recovered 82% of fragments from the 13 homology-detectable genomes (350/427), compared with 66% for HVSeeker (282/427; **Table S37**). Their false-positive rates on bacterial negatives were 11% and 52%, respectively (**Table S1**).

**Figure 5.**
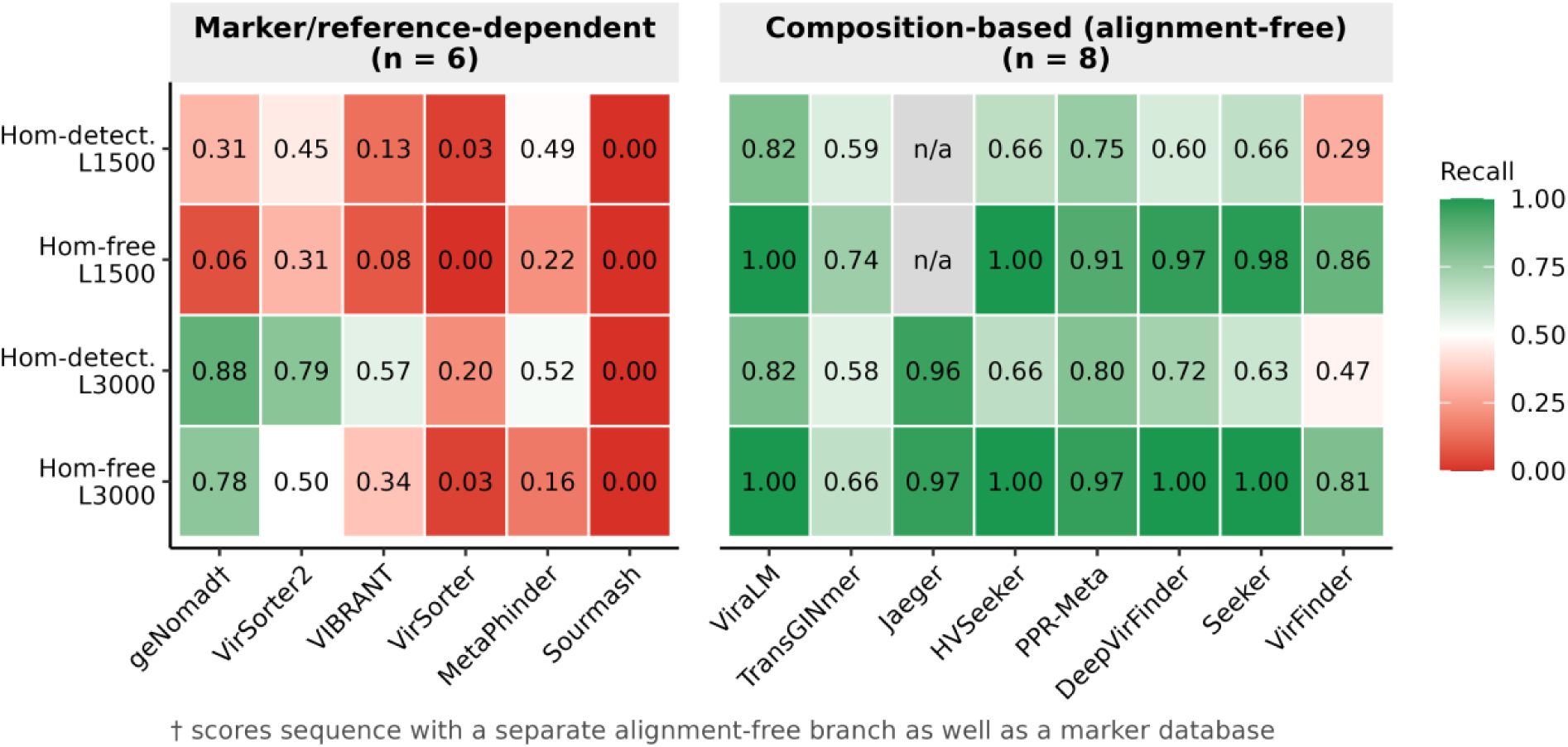
Composition-based tools have higher recall on homology-free vaginal phage fragments at 1,500 bp. Cells show recall for 14 tools across two genome groups and two fragment lengths; numbers and colour indicate recall from 0 (red) to 1 (green). Hom-detect. denotes genomes with detectable reference relatives, and Hom-free denotes genomes without detectable nucleotide or protein homology under the searches used here (Methods). L1500 and L3000 indicate fragment lengths of 1,500 and 3,000 bp. Tools are grouped by dependence on external reference or marker databases (**Table S37**); † marks geNomad, which also uses sequence composition. Grey n/a cells indicate Jaeger’s minimum input length exceeds 1,500 bp. The homology-detectable and homology-free groups contain different genomes, although all tools are evaluated on the same fragments within each group. Sample sizes: 13 homology-detectable genomes (427 and 210 fragments at 1,500 and 3,000 bp, respectively) and two homology-free genomes (65 and 32 fragments).

At 1,500 bp, every marker- or reference-dependent tool recovered ≤ 31% of homology-free fragments (VirSorter2 31%, MetaPhinder 22%, geNomad 6%, VIBRANT 3%, VirSorter and Sourmash 0%). Each of these tools except Sourmash, which made no call in either group, recovered a larger share of homology-detectable fragments (MetaPhinder 49%, VirSorter2 45%, geNomad 31%, VIBRANT 11%, VirSorter 3%; **Table S37**). Every evaluable composition-based tool recovered ≥ 74% (ViraLM and HVSeeker 100%, Seeker 98%, DeepVirFinder 97%, PPR-Meta 91%, VirFinder 86%, TransGINmer 74%). Jaeger requires fragments ≥ 2,048 bp and could not be evaluated at 1,500 bp. At 3,000 bp, geNomad reached 78% recall (25/32), exceeding TransGINmer (66%); VirSorter2 reached 50%, while VIBRANT, MetaPhinder, VirSorter, and Sourmash remained at 25% or below. Jaeger recovered 97% (31/32). At 3,000 bp the two groups overlapped, unlike at 1,500 bp, although recall stayed at or below 50% for five of the six marker- or reference-dependent tools.

A vaginal-specific reference recovered relatives missed by MetaVR. In a cross-check against VMGC database, 12 of 15 panel genomes matched at species level, including both genomes without detectable MetaVR relatives (**Table S25**). Across all 4,263 VMGC vOTUs, 913 (21.4%) had a MetaVR species-level match and 4,212 (98.8%) retained a detectable RefSeq protein homolog (**Table S24**; **Figure S11**).

### Assembly coverage limits recovery, while CST-associated accuracy differences vary among tools

Viral contig recovery increased with read coverage; no viral contigs ≥ 1,500 bp were recovered below 5× coverage (**Figure 6A**; **Figures S6, S18**; **Tables S8, S10**). At 10× coverage, MCC was generally lower in the four CST-IV-B backgrounds than in the four CST-I backgrounds.

**Figure 6.**
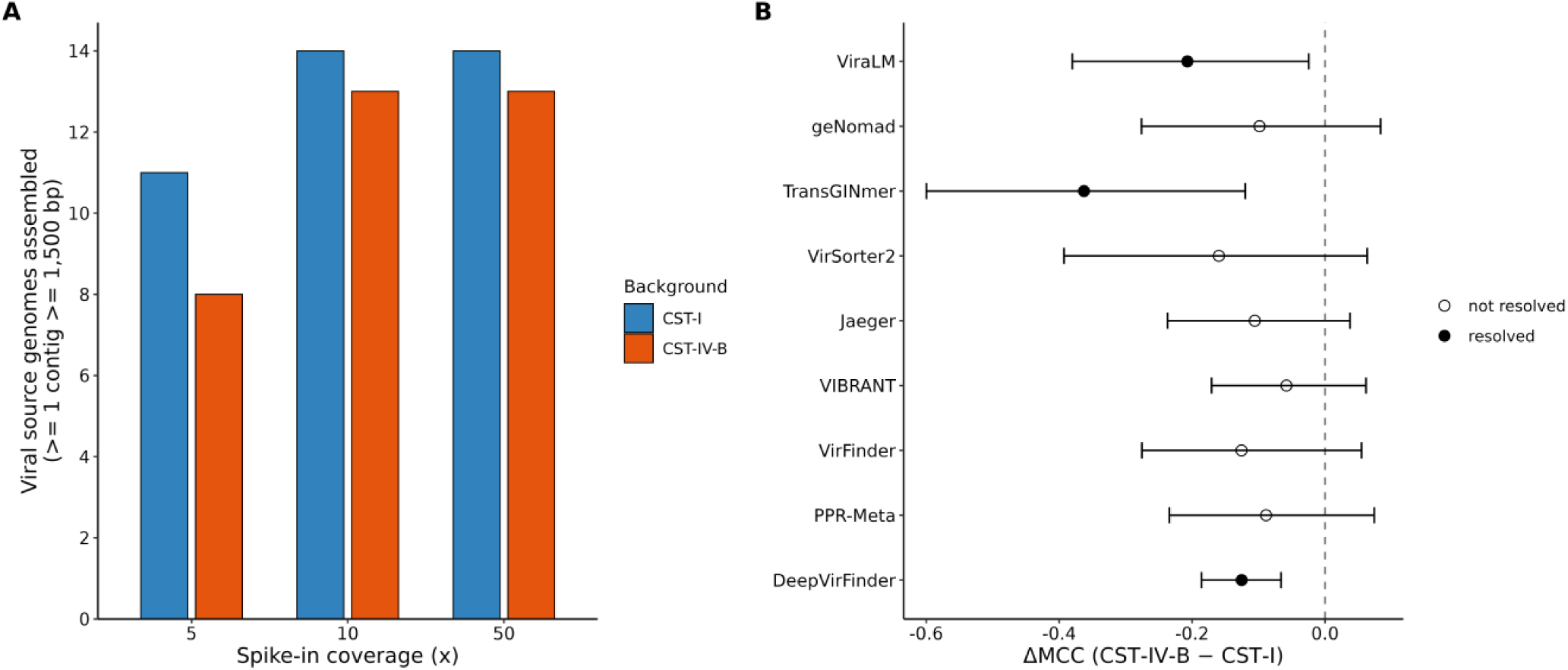
Viral contig assembly increases with coverage, while CST-associated MCC differences vary among tools. (**A**) Number of the 14 spiked viral genomes recovered as at least one contig ≥ 1,500 bp at 5×, 10× and 50× coverage in one CST-I and one CST-IV-B background. No qualifying viral contigs assembled below 5×. (**B**) ΔMCC (CST-IV-B minus CST-I) at 10× coverage for nine tools; all 14 are shown in **Figure S12**. Intervals are background-level bootstrap 95% confidence intervals. Filled points indicate intervals excluding zero; open points indicate intervals spanning zero (dashed line). Native-phage homologs were excluded.

Background-level bootstrap 95% confidence intervals for the difference excluded zero for five tools showing a decrease (ViraLM, TransGINmer, VirFinder, PPR-Meta and DeepVirFinder) and for two showing an increase (Seeker and HVSeeker; **Figure 6B**; **Table S9**). MCC also decreased for geNomad (ΔMCC −0.152) and VirSorter2 (−0.128), but both confidence intervals included zero (**Table S9**). Results for all 14 tools and individual backgrounds are shown in **Figures S12 and S19**.

Mean precision was lower in the CST-IV-B backgrounds for 11 of the 13 tools that made viral calls, including geNomad (0.90 to 0.77), ViraLM (0.83 to 0.63) and VirSorter2 (0.85 to 0.71), while recall changed little (0.80 to 0.81, 0.96 to 0.97 and 0.61 to 0.63, respectively). No tool’s precision difference was resolved (geNomad −0.130, 95% CI −0.373 to +0.038; ViraLM −0.198, −0.519 to +0.087; VirSorter2 −0.133, −0.396 to +0.076; **Table S9**). In a post hoc leave-one-out analysis, omitting one CST-IV-B background, SRR27287964, reduced the differences to −0.013, −0.039 and −0.007, respectively (**Table S9**). That background yielded 2,975 negative contigs and a viral fraction of 3.7%, against 24.7% to 41.9% in the CST-I backgrounds. Of the three tools, only ViraLM had a resolvably higher false-positive rate in CST-IV-B backgrounds (+0.133, 95% CI +0.003 to +0.289; **Table S9**).

### geNomad and VIBRANT lead the multi-evidence benchmark at different operating points

On real assemblies (13,030 contigs ≥ 1,500 bp across 13 samples; **Figure 7A**; **Figure S20**), geNomad and VIBRANT ranked first and second by MCC (0.440 [95% CI 0.414 to 0.464] and 0.439 [0.417 to 0.461]), followed by VirSorter2 (0.382 [0.353 to 0.407]; **Table 1**; **Table S12**). The geNomad–VIBRANT difference was not resolved (paired bootstrap ΔMCC +0.001 [−0.024 to +0.026]), whereas geNomad exceeded VirSorter2 (+0.058 [+0.034 to +0.081]; **Table S27**). The two leaders reached similar MCC by different routes. VIBRANT made 8 false-positive calls among 11,577 negatives (precision 0.975) but recovered 22% of viral contigs, whereas geNomad recovered 33% with 201 false positives (precision 0.703; **Table S12**). ViraLM declined to rank 7 (0.224 [0.203 to 0.246]) on real metagenomes, while Jaeger achieved the highest AUPRC (0.420 [0.391 to 0.445]). Removing the MetaVR nucleotide-homology line, whose construction could favour geNomad (E1b), lowered geNomad’s MCC from 0.440 to 0.402 and placed VIBRANT first (0.448); VIBRANT also ranked first, and geNomad second, in the other two evidence-line checks (**Table S11**). For the read-level RCA signal (evidence line E5), RCA-validated recall spanned 0 to 55% on phage contigs and 0 to 96% on eukaryote-infecting virus contigs, and exceeded phage recall for 9 of the 12 tools returning any positive call (**Figure S7**; **Table S3**).

**Figure 7.**
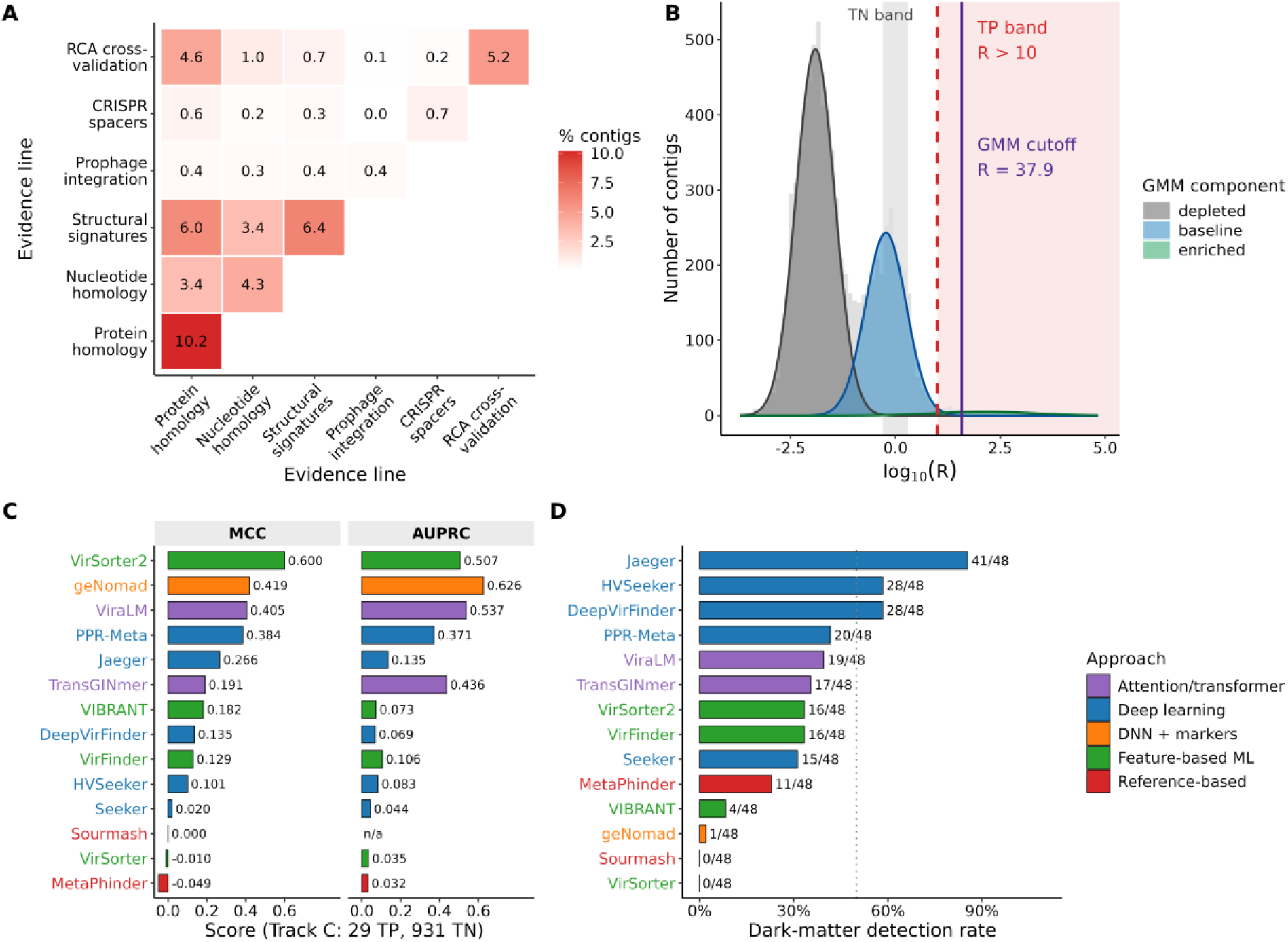
Classification performance and candidate dark-matter recovery on real assemblies. (**A**) Evidence co-occurrence in the multi-evidence benchmark. Cells show the percentage of contigs carrying each evidence category (diagonal) or both categories (off-diagonal); E1 homology is split into protein (E1a) and nucleotide (E1b). (**B**) Distribution of the library-size-normalised RCA enrichment ratio R on a log₁₀ scale. Shading marks the positive enrichment region (R > 10, red) and negative band (0.5 < R < 2, grey); classification also requires annotation evidence (Methods). Curves show the three fitted Gaussian-mixture components; the solid line marks the alternative positive cutoff, R = 37.9 (**Table S23**). (**C**) Track C MCC and AUPRC for all 14 tools. Sourmash’s AUPRC is unavailable because it returns no ranked scores; confidence intervals are in **Table S35**. (**D**) Fraction of 48 candidate dark-matter contigs detected by each tool; the dotted line marks 50%. Candidates have RCA enrichment and no detectable nucleotide or protein homology under the searches used here. Sample sizes across 13 samples: 13,030 contigs (**A**), 8,164 with finite R > 0 (**B**), 960 benchmark contigs (**C**) and 48 candidates (**D**).

**Table 1.** geNomad ranks first or second in every track; VIBRANT reaches the top two only on real shotgun assemblies. Per-tool Matthews correlation coefficient (MCC) across Track A (1,500 bp), Track C, the multi-evidence benchmark, and the MiTCH external validation for the five decision-relevant tools (geNomad, ViraLM, VirSorter2, VIBRANT, Jaeger). Jaeger is undefined at 1,500 bp (2,048-bp floor).

| Tool | Track A MCC<br>(1,500 bp) | Track C<br>MCC | Multi-evidence<br>MCC | MiTCH MCC |
| --- | --- | --- | --- | --- |
| geNomad | 0.651 | 0.419 | 0.440 | 0.457 |
| ViraLM | 0.536 | 0.405 | 0.224 | 0.225 |
| VirSorter2 | 0.283 | 0.599 | 0.382 | 0.308 |
| **VIBRANT** | 0.172 | 0.210 | 0.439 | 0.568 |
| **Jaeger** | N/A (min 2,048 bp) | 0.266 | 0.292 | 0.350 |

Tool errors were not distributed evenly across the negative classes. On the 2,833 negatives grounded by a Kraken2 human assignment, the false-positive rate spanned 0.0% (VIBRANT) to 95.5% (TransGINmer). The ordering differed from that on bacterial negatives. TransGINmer misclassified 95.5% of human contigs as viral against 38.9% of bacterial or archaeal contigs, whereas Jaeger misclassified 0.9% against 20.1%. The 80 human-assigned Track C contigs in the background enrichment band reproduced this ordering (**Table S38**). These contigs carry Kraken2 assignments rather than verified human sequence.

### Marker-dependent tools show higher but unresolved recall on prophage contigs

Marker-dependent tools had higher mean recall on prophage contigs, but the difference was not resolved. To address Q4, recall was compared between tools using viral gene markers and tools using other sequence features on the Tier 1 prophage set (Methods). On the 49 prophage contigs supported by ≥ 3 evidence lines (Tier 1), mean recall was 91.3% for marker-dependent and 77.0% for sequence-based tools (4 and 7 tools; exact permutation Mann–Whitney p = 0.097), with broad variation within both groups (75.5–98.0% and 38.8–91.8%). All 49 carried homology, CheckV and Phigaro support, so the set is restricted to prophages these methods already recognise (Supplementary Methods). With four and seven tools, this test could only have resolved near-complete separation between the groups (smallest attainable two-sided p = 0.006). On the 1,342 free-phage contigs, recall was lower for the marker-dependent tools (mean 21.8% versus 37.0%; p = 0.073; **Table S13**).

### VirSorter2 leads Track C, while tool rankings are broadly concordant across evaluation contexts

On Track C (29 true positives, 931 true negatives; **Figure S21**), VirSorter2 achieved the highest MCC (0.599 [95% CI 0.478 to 0.706]), followed by geNomad (0.419 [0.287 to 0.540]) and ViraLM (0.405 [0.300 to 0.500]; **Figure 7C**; **Table 1**). Paired bootstrap differences excluded zero for VirSorter2 versus geNomad (ΔMCC +0.180 [+0.069 to +0.305]) and versus ViraLM (+0.194 [+0.088 to +0.310]). The geNomad–ViraLM difference included zero (+0.014 [−0.093 to +0.110]; **Table S27**). Track C contigs were longer than Track A fragments (pooled N50 22,105 bp; **Figure S8**). Across ten pairwise comparisons between five evaluation contexts, rankings were broadly concordant (Spearman ρ = 0.51 to 0.91, median 0.74; 9 of 10 significant; **Table S22**). The weakest correlations involved the multi-evidence benchmark compared with Track A at 1,500 bp (ρ = 0.506) and Track C (ρ = 0.596). The Track C ranking was robust to the grid of enrichment-ratio cutoffs defining its positives (*R*; Methods) and to Gaussian-mixture relabelling (the latter at ρ = 0.94; Supplementary Methods; **Tables S18, S23**; **Figure 7B**; **Figure S10**).

### Jaeger detects the most candidate dark-matter contigs

Among 48 contigs enriched by RCA and novel at both the nucleotide and protein levels (median 3.3 kb; **Table S14**), per-tool detection spanned more than an order of magnitude, from 41 of 48 (85%) for Jaeger to 1 of 48 (2%) for marker-weighted geNomad and zero for VirSorter and Sourmash (**Table S15**; **Figure 7D**). VIBRANT, the other leader on real assemblies, detected 4 of 48 (8%). These contigs had no detectable nucleotide or protein relatives, whereas 13 of the 15 genomes in the Track A ANI-novel panel retained detectable protein relatives. Against the VMGC catalogue [14], 47 of 48 dark-matter contigs had no detectable homology (**Table S25**). Lowering geNomad’s default threshold score from 0.7 to 0.30 increased recovery from one to three contigs in total (**Table S17**).

### The tool ranking replicates in an independent validation cohort

The tool ranking replicated in the independent cohort. A matched four-evidence benchmark was applied to 30 MiTCH shotgun metagenomes (Methods), and the per-tool MCC ranking replicated the primary ranking (Spearman ρ = 0.952, 95% CI 0.921 to 0.960; **Figure 8A**). VIBRANT ranked first in MiTCH (pooled MCC 0.568, 95% CI 0.546 to 0.587), ahead of geNomad (0.457 [0.394 to 0.503]; paired ΔMCC +0.111 [+0.070 to +0.166]), Jaeger (0.350 [0.317 to 0.381]) and VirSorter2 (0.308 [0.239 to 0.368]; **Table S27**). In the matched four-evidence primary benchmark the two leaders were nearly tied (0.574 and 0.571). The weak cluster (Seeker, Sourmash, TransGINmer) again ranked last (**Table S27**; **Figure S13**). Absolute accuracy was lower in the validation cohort, with MCC below the matched four-evidence primary benchmark for 12 of 14 tools (median ΔMCC −0.091; geNomad 0.457 versus 0.571; **Table S27**). CST-associated differences depended on how MCC was summarised. In the observational contrast, pooled MCC was higher in high-diversity CST-IV samples (n = 19) than in low-diversity CST-I/III/V samples (n = 11) for several top tools (geNomad ΔMCC +0.176, VirSorter2 +0.155, ViraLM +0.116, all CIs above zero), whereas the differences for VIBRANT (−0.051) and Jaeger (−0.070) were not resolved (CIs spanning zero; **Figure 8B**; **Table S31**). The CST-IV contig sets, however, carried a larger labelled viral fraction (6.4% versus 2.0%), and MCC depends on class prevalence. When samples were weighted equally, or when each tool’s CST-IV sensitivity and false-positive rate were evaluated at the CST-I/III/V viral fraction, the three positive differences were no longer resolved (prevalence-matched ΔMCC +0.011 to +0.031) and VIBRANT’s remained unresolved, whereas Jaeger’s and MetaPhinder’s became negative (−0.195, 95% CI −0.274 to −0.073, and −0.100, −0.173 to −0.041; **Table S39**). Across samples, bacterial Shannon diversity correlated with the number of labelled viral contigs (Spearman ρ = 0.70, n = 30; ρ = 0.90 within CST-IV-B, n = 18) and with the labelled viral fraction (ρ = 0.62; **Table S40B**), and per-sample MCC for geNomad, VirSorter2 and ViraLM increased with viral fraction (ρ = 0.58 to 0.74), whereas VIBRANT’s association was weaker and not resolved (ρ = 0.34; **Table S40**). Within CSTs, VIBRANT led in four of the five CST groups, including CST-IV-B (n = 18) and CST-I (n = 6), and MetaPhinder in the single CST-V sample (**Figure S22**; **Table S28**; by metagenomic CST in **Table S29**), but these groups were small and reported descriptively. Per-sample MCC values for all 14 tools are in **Table S30**.

**Figure 8.**
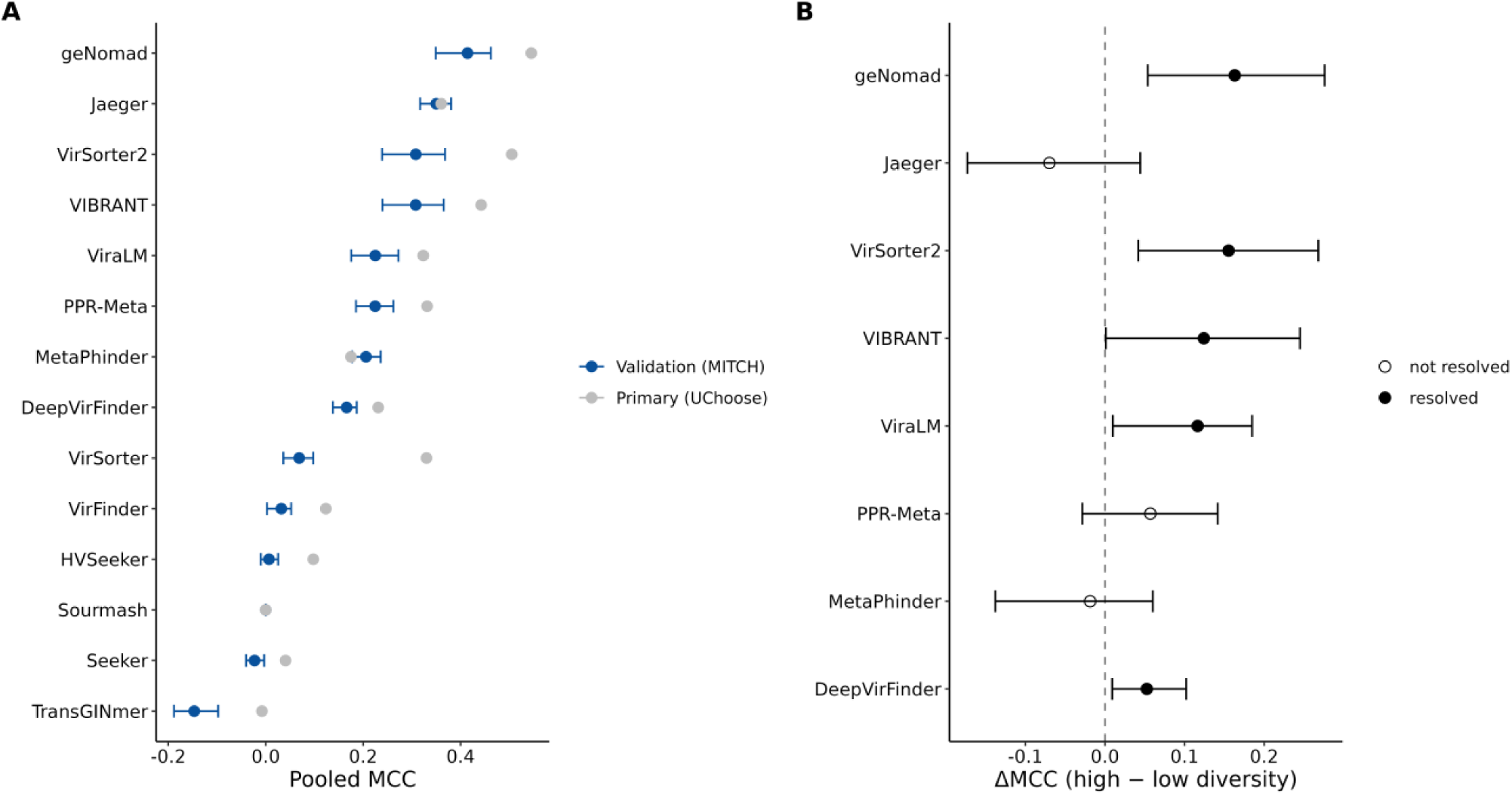
The tool ranking replicates in the independent cohort, while pooled MCC differs by community state type. (**A**) Per-tool pooled MCC in the external validation cohort (MiTCH), with sample-block bootstrap 95% confidence intervals drawn on the validation series only, alongside the matched four-evidence primary (UChoose) ranking, plotted without intervals. Where the two series coincide, as for Sourmash, the grey primary point overplots the blue validation point. The two correlate at Spearman ρ = 0.952. (**B**) Difference in pooled MCC for eight representative tools between high-diversity CST-IV (n = 19) and low-diversity CST-I/III/V (n = 11) samples, with sample-block bootstrap 95% confidence intervals. CST-IV contigs carry a larger labelled viral fraction; equal-weight and prevalence-matched differences are in **Table S39**. The dashed vertical line marks ΔMCC = 0; filled points (legend: CI excludes 0) are intervals excluding zero, open points (CI includes 0) those spanning it. n = 30 samples.

### Computational requirements vary widely across tools

Per-tool runtime at 1,500 bp spanned 206-fold, from 28 seconds (TransGINmer) to 1.6 hours (Sourmash), and peak memory spanned 310-fold, from 61 MB (VirSorter) to 18.5 GB (geNomad, which loads its marker database; **Table S16**; Jaeger, fastest where it runs, undefined below 2,048 bp). No significant association between runtime and MCC was detected across the 13 tools evaluable at this length (Spearman ρ = −0.16, p = 0.61), and 10 of the 13 were both slower and less accurate than TransGINmer, ViraLM, or geNomad (**Figure 9A**). Sourmash spent 1.6 hours to reach MCC 0.000, returning no positive call at any of the six Track A fragment lengths (**Table S7**), and DeepVirFinder 1.4 hours to reach 0.152, against ViraLM’s 0.536 in 43 seconds.

**Figure 9.**
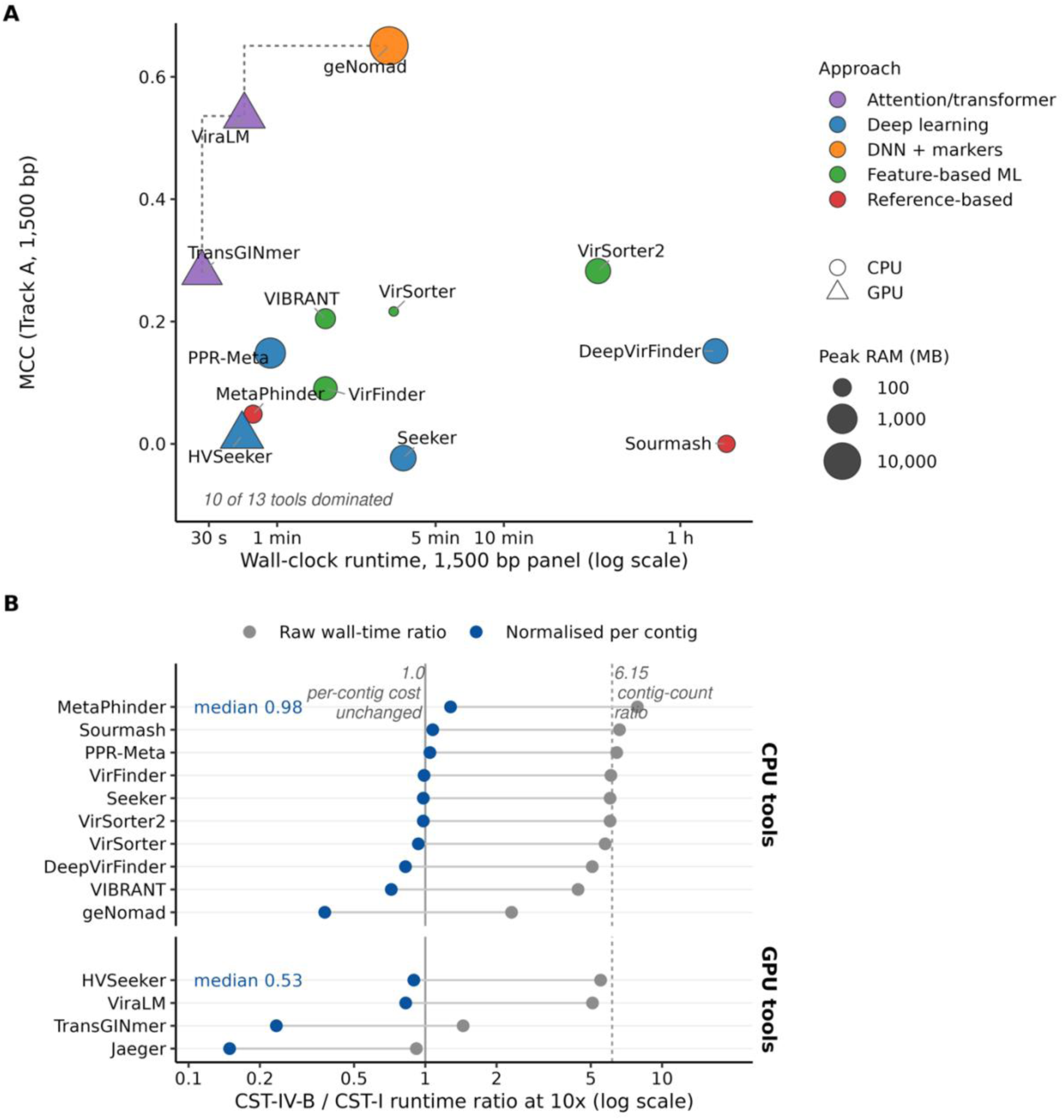
Runtime and memory requirements vary across tools. **A**) Wall-clock runtime versus MCC for 13 tools at 1,500 bp. Runtime and memory size scales are logarithmic; point area represents peak resident memory, colour identifies methodological approach and shape identifies processor. The dashed frontier connects tools that no other tool outperforms on both runtime and MCC. Jaeger is excluded because its minimum input length exceeds 1,500 bp. (**B**) CST-IV-B:CST-I runtime ratios at 10× coverage, before (grey) and after (blue) contig-count normalisation. The solid line marks a ratio of 1; the dashed line marks the contig-count ratio, 6.15 (5,149/837). CPU and GPU panels show median normalised ratios. Timings are single measurements on AMD Zen4 CPUs and NVIDIA L40s GPUs under the documented settings (Methods; **Table S16**). Sample sizes: 13 tools (**A**), 14 tools (**B**)

For CPU tools, the median of per-tool wall-time ratios between the CST-IV-B and CST-I co-assemblies was 6.03 (range 2.31–7.86). The CST-IV-B assembly contained 6.15-fold more contigs (5,149 versus 837; **Table S8**). After normalisation per contig, the median CPU-tool ratio was 0.98 (range 0.38–1.28). Across all 14 tools, the lowest normalised ratios were Jaeger 0.15, TransGINmer 0.23, and geNomad 0.38 (**Figure 9B**). Combining cross-track MCC, recall on eukaryote-infecting viruses and on ANI-novel genomes, community-state performance, and runtime, we derived tool recommendations stratified by research goal, contig length, and resources (**Table 2**).

**Table 2.** The recommended tool depends on the research question, contig length and available compute. Tools are grouped as recommended or complementary, with the use each suits best, the benchmark evidence supporting it, and its compute and minimum-length requirements. Recommendations apply at default score cutoffs. MCC, Matthews correlation coefficient; FPR, false-positive rate; ORF, open reading frame; VLP, virus-like particle; MiTCH, external validation cohort.

|  |  |  |  |  |
| --- | --- | --- | --- | --- |
| **Recommended** | geNomad | High-confidence identification across contig types | Highest Track A MCC; first or second by MCC in every track | CPU (18.5 GB) |
| **Recommended** | VIBRANT | High-precision phage catalogues from assembled contigs | Highest MiTCH MCC; almost no false-positive calls on real assemblies; 98.0% prophage recall | CPU; min 1,000 bp and $\geq 4$ ORFs |
| **Recommended** | ViraLM | Short contigs, discovery and eukaryote-infecting virus detection | Only tool with 100% eukaryotic recall; robust on novel genomes | GPU (2.1 GB); min 500 bp |
| **Recommended** | Jaeger | Novel phage discovery (paired with geNomad) | Recovered 41/48 dark-matter contigs; fastest on large inputs | GPU (4.6 GB); min 2,048 bp |
| **Recommended** | VirSorter2 | VLP-enriched contigs | Highest Track C MCC (0.599); on VLP-enriched co-assemblies | CPU |
| **Complementary** | PPR-Meta | Phage detection on long contigs | Ranked 4th on Track C; three-class model controls FPR | CPU |
| **Complementary** | DeepVirFinder | Broad phage screening | 58% dark-matter detection; high *Lactobacillus*-phage recall | CPU |

## Discussion

Our benchmark shows that tool choice affects both viral recovery and false-positive calls in vaginal metagenomes. Across 14 tools, geNomad ranked first or second by MCC in every ranked evaluation. VIBRANT achieved similar MCC on the primary shotgun assemblies while recovering fewer viral contigs and making very few false-positive calls. It also ranked first in the independent cohort. Jaeger flagged the most candidate dark-matter contigs. Earlier benchmarks, run on simulated or non-vaginal data with partly overlapping tool panels, favoured other tools [38, 40]. Our results extend those evaluations to vaginal metagenomes and show why tool selection should reflect the study objective. Their applicability to other viral groups and sequencing approaches remains uncertain.

Tool rankings agreed across evaluations even when absolute performance changed. Rankings correlated across tracks with different labelling assumptions (Spearman ρ = 0.51–0.91, Table S22). Agreement was stronger between the primary and independent cohorts (ρ = 0.952). Nevertheless, 12 of 14 tools had lower MCC in the independent cohort than in the matched primary benchmark. Within the primary multi-evidence benchmark, the leading tool also depended on the evidence used to label contigs. Removing MetaVR evidence placed VIBRANT first. Because MetaVR was built using geNomad [13] including this evidence could favour geNomad. Correlated fragments and repeated samples further limit independence and may make uncertainty intervals too narrow. The ranking agreement therefore supports consistency across the evaluated datasets, while leaving the exact order of the leading tools uncertain.

VirSorter2’s leading performance on VLP-enriched contigs requires confirmation in a larger dataset. Track C contained 29 labelled viral contigs from 13 samples collected from 11 participants. Its labels also depended on library preparation. We used enrichment relative to matched shotgun libraries from the UChoose cohort [42], whose virome filtrates were not DNase-treated. Rolling-circle amplification favours circular templates [43, 44], so these labels may over-represent circular viral genomes. Residual free DNA could also reduce enrichment ratios for viral contigs. Non-viral circular elements, including plasmids, could enter the candidate set. Validation against independently confirmed viral contigs would clarify whether this advantage persists.

Sparse reference coverage of vaginal viruses is the likely constraint behind the largest differences in tool accuracy. Only 21.4% of 4,263 species-level units in the vaginal-specific VMGC catalogue [14] matched a MetaVR species ([13]; **Table S24**). Each evaluated composition-based tool recovered at least 74% of 1,500-bp fragments from the two phage genomes without detectable reference homologues. Marker- or reference-dependent tools recovered at most 31%. For geNomad, recall was 6%, compared with 31% for fragments from the 13 genomes with detectable homologues (**Table S37**). These results do not isolate reference coverage as the cause. The two genomes may be unusually recognisable by composition. Part of geNomad’s underperformance reflects a marker requirement in its reporting rule, which withholds calls below 2,500 bp that lack a virus hallmark gene. Higher recovery by composition-based tools must also be weighed against false-positive calls, a trade-off reported in an earlier benchmark [39].

Of 48 dark-matter candidates, sequences without detectable homology to reference viruses [45, 46], Jaeger detected 41 and geNomad one. These candidates are unconfirmed and include no negatives. They therefore show which tools flag novel sequence, not which flag it correctly. No contig was validated experimentally, and novelty here is relative to the resources searched. Nondetection of a phage group in vaginal data should not be read as biological absence.

Detection also declines unevenly across the phages whose bacterial hosts are of clinical interest. Every tool that recovered *Gardnerella* or *Lactobacillus* phage fragments recovered fewer *Gardnerella* fragments, at 1,500 bp and at 3,000 bp (**Figure 4**). The comparison rests on one *Gardnerella* and two *Lactobacillus* phage genomes. Composition-based tools showed the same gap, so reference coverage is unlikely to be its only cause. Published associations between phage communities and BV rest on homology-based identification [8, 34]. They may therefore under-represent phages of BV-associated bacteria, with effects of unknown direction. Prophages that carry recognisable viral genes are a partial exception. Nine of the 14 tools recovered at least 40 of the 49 highest-confidence prophage contigs, with no resolved advantage for marker-dependent tools. This set contains only prophages that homology searches, CheckV and Phigaro already recognise, so it likely overstates prophage recall in general. Their detection is better served than their classification. In an earlier analysis of the same parent cohort, no viral family or host group could be assigned to the putative *G. vaginalis* prophages identified there [30].

Comparisons between CST-IV and *Lactobacillus*-dominated communities are not measurement-neutral. In the independent cohort, CST-IV assemblies carried about three times the labelled viral fraction of CST-I/III/V assemblies (6.4% versus 2.0%). Bacterial diversity tracked that fraction (Spearman ρ = 0.62; **Table S40**). At identical error rates, a smaller share of a tool’s calls is then false in CST-IV. Once the viral fraction was matched, the higher pooled MCC of geNomad, VirSorter2 and ViraLM there was no longer resolved. Jaeger and MetaPhinder instead scored lower in CST-IV (**Table S39**). A co-assembly contrast used four CST-I and four CST-IV-B backgrounds, at one coverage and in one CST-IV sub-type. MCC was resolvably lower in CST-IV-B for five tools, including ViraLM, and resolvably higher for Seeker and HVSeeker, but unresolved for geNomad and VirSorter2 (**Table S9**). Its backgrounds ranged from 3.7% to 92.5% viral, so it too cannot separate community type from prevalence (Supplementary Methods). Omitting its 3.7% viral background alone nearly removed the precision differences for geNomad, ViraLM and VirSorter2 (**Table S9**). Operational negatives may also contain unrecognised viruses, more often in CST-IV-B backgrounds, which would inflate apparent false-positive rates there. Because Amsel-diagnosed BV maps onto only some metagenomic community state types [47], these effects may carry over to BV comparisons only in part. Differences in viral richness between BV and non-BV groups can still be produced or masked by detection behaviour. They carry more weight when class prevalence is checked and the result holds across tools. The share of CST-IV rose with cervical lesion stage among 68 women with high-risk HPV [48], so virome comparisons across lesion stages need the same prevalence check.

For eukaryote-infecting viruses, detection reliability differs by virus and by tool. Human papillomavirus (HPV) fragments were recovered by more tools than herpesvirus fragments, including several built for phages. HPV recall at 1,500 bp nonetheless ranged from none to complete across tools. Recall is informative only where a tool also rejects non-viral sequence. Seeker and HVSeeker recovered at least 90% of HPV fragments while labelling more than half of bacterial fragments viral. Phage-oriented tools fail in the opposite direction. VIBRANT recovered no eukaryote-infecting fragment in the panel tested here, so a virome screened that way is a phageome by construction. ViraLM recovered all at 1,500 bp, at an 11.4% bacterial false-positive rate, although unresolved overlap with its training sources qualifies that result. Only tools declaring all-virus scope recovered most herpes simplex virus fragments. Training composition is a plausible contributor. ViraLM was trained on viral genomes spanning six realms [27]. VIBRANT, by contrast, was built for prokaryotic viruses [20]. Scope, architecture and training were not varied independently, however. These viruses have been linked to BV status [8], and their richness to preterm delivery in targeted capture data [49]. Shotgun replication of both associations would therefore be tool-dependent, and a null result hard to distinguish from low recall. Studies of eukaryote-infecting viruses should demonstrate recovery for the specific viral group of interest.

Contig length constrains tool choice in a different way. On 500- and 1,000-bp fragments, attention-based tools outperformed feature-based tools on AUPRC and recall, largely through ViraLM. Their precision was lower than that of the gene-based tools, and geNomad kept the highest MCC at both lengths. Gene-level dependence offers a plausible basis for that gap. VIBRANT skips sequences shorter than 1,000 bp or with fewer than four predicted open reading frames [20]. VirSorter2 scores features computed from predicted genes and is reported to be suboptimal below 3 kb [19]. The k-mer-based VirFinder, which needs no gene calls, was nonetheless also near chance at 500 bp. No viral contig of at least 1,500 bp was recovered below 5× read coverage in the spike-in benchmark (**Figure 6A**). Since short-read assembly bounds contig length, sequencing depth and assembly quality partly determine which tool is appropriate.

The vaginal niche concentrates these detection problems, because its virome is dominated by phages [8] and its bacterial communities by a small number of genera, mainly *Lactobacillus* and *Gardnerella* [32, 33]. Phage profiles in turn track bacterial community composition [8, 34]. Where few hosts dominate, uneven recovery across phage groups acts directly on the comparison of interest rather than averaging out. In this niche, detection behaviour is not a neutral layer over the biology but part of what the biology appears to be.

We therefore recommend matching the tool to the research question rather than adopting a single default (**Table 2**). geNomad suits high-confidence identification and VIBRANT suits phage catalogues where false positives must be minimised. These recommendations apply at default score cutoffs, and the in-sample optima are upper bounds, not transferable settings. CheckV can be coupled with any of these tools to assign quality tiers and remove host regions from proviruses [50]. It was not designed to remove false-positive predictions, however, so catalogue-grade use of a sensitive tool still needs independent quality control. ViraLM’s false-discovery rate, for example, was 6.3-fold higher than geNomad’s at 1,500 bp (0.645 versus 0.103; **Table S34**). Combined workflows, such as pairing Jaeger with geNomad, were not tested here, and the literature does not settle them [39, 40]. Runtime varied 206-fold with no detectable association with MCC, so memory and accelerator access are likely to constrain selection more than speed. These findings support tool selection for the panels and assemblies evaluated here, not inferences about viral identity, activity or health outcomes. Better detection will need vaginal sequences in reference and training databases. For findings on vaginal microbiome-associated outcomes to be comparable across studies, reports will also need to state the tool, version, settings and which output was scored. Long-read assemblies are becoming testable as vaginal Nanopore metagenomes begin to appear [51]. A benchmark pairing VLP-enriched short- and long-read libraries from the same women, with experimental confirmation of candidate contigs, would test whether these rankings hold for the viruses most relevant to these outcomes.

## Materials and Methods

### Study design and data sources

Fourteen viral identification tools were benchmarked using three tracks and a multi-evidence evaluation (**Figure 1**). Track A used controlled fragments of 14 vaginal-associated viral genomes and 6 bacterial negative genomes. Track B co-assembled simulated viral reads with two UChoose backgrounds for a coverage sweep and eight public backgrounds from three cohorts for a 10× diversity contrast between CST-I and CST-IV-B. The multi-evidence benchmark evaluated the 13 UChoose shotgun assemblies, while Track C used the matched shotgun-plus-RCA co-assemblies, labelled by library-size-normalised phi29 enrichment and annotation. An independent set of 30 MiTCH shotgun metagenomes provided external validation.

Thirteen paired samples from 11 UChoose participants among South African adolescents were analysed; UC055 and UC093 each contribute two visits. Each sample has a shotgun metagenome (PRJNA767784; [30]; Illumina NovaSeq 6000) with a matched virome prepared from a 0.2 µm filtrate and amplified by phi29 rolling-circle amplification (PRJNA881266; [42]). CST assignments used VIRGO2 [15] and the VISTA classifier ([52]; extending the mgCST framework of [47]), each metagenomic CST mapped to its traditional CST by species-level dominance (**Table S2**). The 13 samples comprised five CST-I, four CST-III, and four CST-IV (CST-II and CST-V absent). The parent study was approved by the University of Cape Town HREC (REF 801/2014) with informed consent. This is a secondary analysis of deidentified data.

### Read preprocessing and assembly

Reads were trimmed (fastp v1.0.1; [53]) and host-depleted against hg19 with Bowtie2 v2.5.4 (-- *very-sensitive-local*; [54]) and read pairs with both mates unmapped were retained (SAMtools v1.21; [55]). Per-sample metagenomes were assembled with metaSPAdes v4.2.0 ([56]; k-mers *33, 55, 77*) and filtered to ≥ 1,500 bp (**Figure S1**). Shorter Track A fragments were generated separately. Track B and Track C co-assemblies used the same configuration with the modifications below.

### In silico spike-in benchmark (Track A)

The spike-in panel comprised 14 viral genomes (seven vaginal-associated bacteriophages, four high-risk HPV types, torque teno virus 1, and two herpes simplex viruses) and 6 vaginal bacterial type strains as negatives (**Table S33**). Each genome was fragmented into non-overlapping segments at six target lengths (500, 1,000, 1,500, 3,000, 5,000, 10,000 bp), discarding sub-target terminal fragments. At 1,500 bp this produced 421 viral and 6,268 bacterial fragments (1:15). To separate generalisation from training-database memorisation, we also fragmented an ANI-novel panel of 15 vaginal bacteriophage genomes (< 95% ANI to MetaVR; species-level vOTU threshold [57]), split by MetaVR homology into homology-detectable (n = 13) and homology-free (n = 2) subgroups (Supplementary Methods; **Table S25**).

### Assembly-based spike-in benchmark (Track B)

Paired-end Illumina reads (150 bp, NovaSeq error model) were simulated from the viral spike-in genomes (InSilicoSeq v2.0.1; [58]) at six depths (0.1×–50×), mixed with host-removed reads from two backgrounds (CST-I and CST-IV-B), and co-assembled with metaSPAdes (*k-mers 33, 55, 77*; ≥ 1,500 bp). Contigs were classified by origin with minimap2 v2.29 ([59]; asm5, ≥ 95% identity, ≥ 80% query coverage). For the community-diversity contrast, the panel was co-assembled at 10× coverage into eight real backgrounds (four CST-I, four CST-IV-B) from three public cohorts (PRJNA1054643, PRJNA1356845, PRJNA1288683), with backgrounds as the unit of replication. Cohort representation was unequal between the CST groups (Supplementary Methods). Contigs matching excluded native viral homologs were removed before scoring; the remaining background-origin contigs were scored as negative, without establishing that each was nonviral (Supplementary Methods).

### Multi-evidence benchmark for real metagenome assemblies

Each real-metagenome contig (≥ 1,500 bp) was scored on five viral evidence lines (**Figure S14**): homology (E1a, DIAMOND BLASTx against RefSeq viral proteins, [60], and E1b, BLASTn against MetaVR, [13, 61], counted as one line), structural signatures (E2; CheckV, [50]), prophage integration (E3; Phigaro, [62]), CRISPR-spacer targeting (E4; CRISPRCasFinder, [63]), and RCA cross-validation (E5; minimap2, [59]). Contigs were graded by how many lines supported them, whereas negatives required an explicit non-viral signal (a Kraken2 non-viral assignment, [64], or CheckV host genes without viral genes). Per-line thresholds are in **Table S36**, the tier rule and the contig count at every step in **Figure S14**, and the composition of the retained sets in **Figure S15**.

E5 measures RCA read support on shotgun contigs; Track C instead uses a library-size-normalised enrichment ratio on co-assemblies. Evidence-line dependencies, the sensitivity of evidence tiers to E1b coverage estimator, and threshold sensitivity analyses are described in Supplementary Methods (**Tables S11, S19, S26**; **Figures S9, S16**).

### VLP-enrichment benchmark (Track C)

For each matched sample, host-removed shotgun and RCA reads were co-assembled with metaSPAdes (*k-mers 33, 55, 77*, *--only-assembler*) and filtered to ≥ 2,500 bp. The 0.2 µm filtrate was not DNase-treated [42]. Phi29 enrichment was quantified from the paired libraries (Supplementary Methods). Each library was mapped to the master contigs (Bowtie2 v2.5.4), giving the library-size-normalised enrichment ratio,

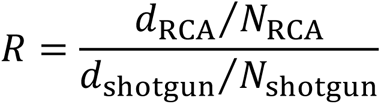

where *d* is the per-contig mean depth and *N* is the total number of mapped reads.

Contigs were annotated (CheckV v1.0.1, Kraken2 v2.17.1) and labelled true positive (R > 10 with a CheckV viral signal), true negative (0.5 < *R* < 2 with a Kraken2 bacterial or archaeal call and no CheckV viral signal), or dark-matter (*R* > 10 with no CheckV viral signal, no CheckV host genes, and no Kraken2 classification). Human reads were depleted before assembly; residual human or other eukaryotic assignments do not qualify as Track C negatives under this rule. Other contigs were excluded (full conjunctive rules in Supplementary Methods). The Track C sets were constructed independently of the ANI-novel panel. Robustness was assessed with an R-cutoff grid and a Gaussian-mixture relabelling (scikit-learn; [65]; Supplementary Methods; **Tables S18, S23**; **Figure S10**).

### External validation cohort

To test out-of-sample generalisation, we applied the benchmark to 30 shotgun metagenomes from an independent Swedish cohort (MiTCH study, [66]), labelled MITCH01–MITCH30. Ground truth used four evidence lines: E1 homology (E1a and E1b counted together), E2 structural signatures, E3 prophage integration, and E4 CRISPR-spacer targeting. E5 was unavailable without paired viromes, so comparisons use the primary cohort’s matched no-E5 benchmark.

VIRGO2 and VISTA supplied the mgCST labels used for sample selection. VALENCIA [67] supplied traditional CST labels for the diversity contrast (**Table S32**). The primary analysis had already mapped mgCSTs to traditional CSTs by dominant species (**Table S2**); MiTCH used a direct VALENCIA assignment. Tools were run once on the merged, sample-prefixed Tier 0–2 contigs, and arm comparisons used a sample-block bootstrap (B = 2,000). The primary arm contrast used pooled MCC (confusion counts summed across samples; **Table S31**). Two sensitivity analyses used the same bootstrap draws: the mean of per-sample MCC, and pooled MCC recomputed with each arm’s sensitivity and false-positive rate held fixed and evaluated at a common labelled viral fraction, either that of CST-I/III/V or that of the whole cohort (**Table S39**). Bacterial Shannon diversity was computed from VIRGO2 taxon abundances and related to each sample’s labelled viral fraction and MCC by Spearman correlation, with permutation p-values (10,000 permutations) and bootstrap 95% confidence intervals (**Table S40**).

The parent MiTCH study was approved by the Swedish Ethical Review Authority (approval 2021-05794-01), and all participants provided written informed consent.

### Tool panel and execution

Published tools were included if they returned a viral score or call for each input sequence, directly or through per-sequence invocation. Sequences a tool did not report were scored as negative. Read-based taxonomic profilers, including BAQLaVa [68], were excluded because their taxon-abundance outputs and library-level detection rules do not provide equivalent per-contig classifications. The 14 tools span five methodological approaches (Reference-based, Feature-based ML, Deep learning, Attention/transformer, DNN + markers) and differ in their published target scopes (**Table S21**). A separate grouping distinguishes six tools that require an external reference or marker database from eight composition-based tools (**Table S37**); this grouping differs from the approach categories in **Table S21**, and hybrid tools, including geNomad, are assigned by their database requirement. Approaches, scopes, training databases, versions, container digests, thresholds, and non-default invocations are in **Table S21**. No tool was trained on IMG/VR v4 itself. geNomad’s corpus included IMG/VR v3 with GenBank and curated viral sources [29]. Operating rules were geNomad virus score ≥ 0.7, applied to whole contigs and to reported proviral regions; ten score-based tools, score ≥ 0.5; Sourmash containment ≥ 0.5; VirSorter categories 1–4; and, for VIBRANT, its final reported viral sequences (‘phages_combined’ output), the single set of confident predictions its workflow produces [20]. DeepVirFinder and VirFinder additionally required p < 0.05. For tools that report excised regions (geNomad, VIBRANT), a contig was scored as viral if any region reported from it was accepted (Supplementary Methods). Five tools were run with non-default settings (**Table S21**), notably VirSorter2 --provirus-off. Provirus detection was also enabled in a control run on sample UC093_V3 (**Table S21**). Tools ran in Singularity/Apptainer containers, CPU tools on AMD Zen4 CPUs and the four GPU tools (HVSeeker, Jaeger, TransGINmer, ViraLM) on NVIDIA L40s 48 GB GPUs.

### Evaluation metrics and statistical analysis

Recall (sensitivity) was the fraction of viral sequences recovered. Precision was the fraction of viral calls that were correct. MCC incorporates true positives, true negatives, false positives and false negatives. MCC was the primary ranking metric because it remains informative under class imbalance, and non-viral sequences predominated in our test sets [41]. F1 and the threshold-independent AUPRC were also computed. Performance was stratified by contig length, virus category (free phage, prophage, eukaryote-infecting virus), methodological approach, and CST. The 95% confidence intervals were estimated by nonparametric bootstrap over contigs (1,000 resamples, seed 42, percentile interval), applied identically to Track A, the multi-evidence benchmark and Track C, with intervals for all 14 tools tabulated per track (**Tables S4, S12, S35**). Head-to-head differences between named tools were tested by paired bootstrap of ΔMCC on the same contigs (2,000 resamples). Track A tool heterogeneity was assessed with Cochran’s Q on the per-fragment correctness matrix, and 91 pairwise McNemar tests were performed per length with Benjamini–Hochberg correction (q < 0.05; **Figure S3**). Each question (Q1–Q4) used the metric appropriate to it (Supplementary Methods). Viral hallmark genes detectable by hidden Markov models can persist within bacterial sequence. Marker-dependent tools might therefore have an advantage on prophage contigs, which Q4 tested. The Q4 contrast compared four marker-dependent tools (geNomad, VIBRANT, VirSorter, VirSorter2) with seven sequence-based tools, excluding Jaeger and Seeker for length incompatibility and Sourmash for degenerate output (no positive call on any prophage or free-phage contig; **Table S13**), and summarised each group’s recall by its mean. Recall distributions were compared by exact two-sided Mann– Whitney test that enumerated all group assignments using mid-ranks, so that tied recalls were retained, with p taken as twice the smaller tail and the tool as the unit of analysis. Cross-track consistency was assessed by Spearman correlation across five contexts: Track A at 1,500 and 3,000 bp, Track B at 10× in the CST-I coverage-sweep background, the multi-evidence benchmark, and Track C. All 14 tools were included, with Jaeger’s unavailable 1,500-bp MCC coded as zero in this analysis (**Table S22**). The Track B CST and external-validation diversity contrasts used background- and sample-level bootstraps (B = 2,000), distinct from the contig-level bootstrap above. Out-of-sample replication was the Spearman correlation between primary-cohort and validation per-tool MCC rankings.

## Data availability

Raw UChoose sequencing data are available from the NCBI Sequence Read Archive (SRA) under accessions PRJNA767784 (shotgun metagenomes; [30]) and PRJNA881266 (RCA-enriched viromes; [42]). The eight primary Track B diversity backgrounds came from PRJNA1054643, PRJNA1356845, and PRJNA1288683. Of the 30 MiTCH samples analysed here (MITCH01–MITCH30), 19 are available in the European Nucleotide Archive (ENA) under accession PRJEB108308 [66]; the remaining 11 are available upon request. The analysis pipeline and benchmarking code (MIT license) are available at https://github.com/ruqse/vaginal_virbench. Supplementary Methods and Figures S1–S22 are provided in Supplementary_Information.pdf. Tables S1–S41 are provided in supplementary_tables.xlsx, with one table per worksheet.

## Acknowledgments

This work was supported by the SciLifeLab and Wallenberg Data-Driven Life Science (DDLS) program (grant KAW 2020.0239 to L.W.H.). Computational resources were provided by the National Academic Infrastructure for Supercomputing in Sweden (NAISS), funded by the Swedish Research Council. We thank the UChoose study team for generating and depositing the paired shotgun and virome sequencing data analysed here, and Hana Shabana and Kenny Rodriguez-Wallberg for the MiTCH validation-cohort sequencing data.

## Author contributions

F.D. contributed to conceptualization, methodology, software development, formal analysis, investigation, visualization, and writing of the original draft. L.W.H. contributed to conceptualization, funding acquisition, supervision, and review and editing of the manuscript. A.U.H. contributed to UChoose data curation and review and editing of the manuscript. H.B.J. contributed to UChoose data curation and review and editing of the manuscript.

