## Supplementary Methods for "Benchmarking virus identification tools on vaginal metagenomes"

This document contains Supplementary Methods, Supplementary Figures S1 to S22, and legends for Supplementary Tables S1 to S41. The table data are in the accompanying workbook `supplementary_tables.xlsx`, one sheet per table.

### Supplementary Methods

#### Track C enrichment filter: derivation and caveats

The published protocol for the RCA virome libraries (PRJNA881266; Happel et al., 2023) reports 0.2 µm filtration but no DNase step. Co-extracted bacterial DNA could therefore be sequenced alongside virus-like-particle DNA. A binary "present in the virome library" rule could mislabel co-extracted bacterial contigs as viral. Phi29 can preferentially amplify circular templates (Kim et al., 2008; Kim and Bae, 2011). We used paired-library enrichment as one component of the labelling rule, combined with annotation evidence; amplification response alone does not establish a contig's origin or template topology.

The library-size-normalised enrichment ratio $R$ (Methods, "VLP-enrichment benchmark") quantifies this enrichment: $R > 1$ indicates enrichment in the RCA fraction, $R$ close to 1 indicates no enrichment, and $R < 1$ indicates depletion. The distribution of $\log_{10}(R)$ is multimodal, with a baseline component near $R = 1$ and an enriched component at $R$ well above 10. Conservative round-number cutoffs were placed in the valley between components rather than fitting an explicit valley-finder. The Gaussian-mixture robustness check gave similar tool rankings to these cutoffs (Spearman rho = 0.94; Methods; Table S23; Fig. 7B).

Reads were not deduplicated. Library-size normalisation adjusts for the total mapped reads in each library but does not correct template-specific amplification bias.

Phi29 can also amplify bacterial plasmids and other circular extrachromosomal elements, so high $R$ alone does not establish viral origin. Every Track C true positive therefore requires $R > 10$ together with a CheckV viral signal, and enriched contigs with non-viral annotation but no CheckV viral signal are excluded. Viruses with linear genomes may be under-represented in Track C. The multi-evidence benchmark provides a complementary assessment that does not require strong RCA enrichment for every positive contig.

#### Track C ground-truth labelling: full classification rules

Pooled, sample-prefixed master contigs ( ≥ 2,500 bp) were annotated with CheckV v1.0.1 and Kraken2 v2.17.1 (Standard-16 database) and labelled by the conjunctive rules summarised in the main-text Methods. Contigs were classified as **true positives** if $R > 10$ with an independent CheckV viral signal (viral_genes ≥ 1, quality ≥ Medium-quality, or provirus = Yes); as **true negatives** if $0.5 < R < 2$ with Kraken2 Bacteria or Archaea classification and no CheckV viral signal; and as **dark-matter contigs** if $R > 10$ with no CheckV viral signal, no CheckV host genes, and no Kraken2 classification. Moderate-enrichment ($2 \le R \le 10$), depleted ($R < 0.5$), and signal-conflicted contigs (for example, enriched contigs with host genes but no viral genes) were excluded from benchmarking. This yielded 29 true positives and 931 true negatives (class ratio 1:32) and 48 dark-matter contigs. Throughout, "dark-matter contigs" refers only to these 48 RCA-enriched contigs and not to the ANI-novel panel (Track A); the two sets are disjoint by construction (zero pairwise BLASTn hits at ≥ 80% identity and ≥ 50% query coverage).

**Track C robustness checks: grid and mixture-model definitions**

Having defined the Track C ground truth, we tested whether the tool rankings survive the threshold choices used to construct it through two robustness checks. First, we re-scored all 14 tools by MCC under a 3 × 3 grid of alternative cutoffs (TP at $R > 5$, $> 10$, or $> 20$; TN bands 0.33–3.0, 0.5–2.0, or 0.8–1.25; nine combinations in total), recomputing the per-tool confusion matrix at default classification thresholds in each cell and ranking tools by MCC. Second, as a methodologically distinct alternative to the round-number cutoffs, we fit a Gaussian mixture model on $\log_{10}(R)$ (scikit-learn v1.5 `GaussianMixture`, full covariance, n_init = 10, random_state = 0; Pedregosa et al., 2011) over the 8,164 contigs with $R > 0$, finite, and length ≥ 2,500 bp, selecting the component count by Bayesian Information Criterion (3 components; per-component parameters and fitted-model BIC in Table S23). Replacing the round-number $R$ thresholds with posterior-probability cutoffs of 0.95 (TP candidate at P(enriched | R) > 0.95; TN candidate at P(baseline | R) > 0.95) while keeping the conjunctive CheckV and Kraken2 rules unchanged yielded a GMM-derived TP / TN / dark-matter set (Table S23; Fig. 7B) that we re-scored against all 14 tools and compared to the primary ranking by Spearman rank correlation ($\rho = 0.94$, $n = 14$).

**Evidence-5 threshold robustness (multi-evidence benchmark)**

Separately from the Track C ground truth, we evaluated whether the RCA-supported recall ranking is stable to the read-level thresholds that define an RCA-positive contig in Evidence line 5 (E5), under a 4 × 4 grid (min_breadth_pct ∈ {25, 45, 70, 95} × min_read_pairs ∈ {2, 7, 10, 41}; sixteen combinations in total, set as rounded quantiles of the pooled distribution and spanning the production values of 70% breadth and 10 read pairs). For each cell we re-classified the shotgun contigs and recomputed per-tool RCA-supported recall at default classification thresholds, ranking tools by recall (Table S19; Fig. S9). This grid perturbs E5 of the multi-evidence benchmark rather than the Track C ground truth, and the recall it reports is agreement with RCA read support, not recall against the Track C labels; the two therefore rank tools differently.

**Track A ANI-novel panel and Track B origin assignment**

The ANI-novel panel comprised 15 vaginal phage genomes from unpublished vaginal metagenome assemblies, each sharing less than 95% average nucleotide identity with any of the 24.4 million UViGs in MetaVR (the species-level vOTU threshold; Roux et al., 2019). It was divided by MetaVR homology into a homology-detectable group (n = 13 genomes retaining relatives at 84.3–94.9% BLASTn nucleotide identity to MetaVR) and a homology-free group (n = 2 genomes with no MetaVR nucleotide or RefSeq viral protein hit under the initial searches); recall on the two groups is compared at 1,500 and 3,000 bp in Fig. 5. The two strata comprise different genomes, while every tool was evaluated on the same fragments within a stratum. Both genomes without initial reference matches had species-level VMGC matches in the post-hoc cross-check (Table S25). This panel tests novelty relative to the named references and does not demonstrate exclusion from all tool training datasets. For the two-background Track B coverage sweep, spike-in genomes were pre-screened against the backgrounds with BLASTn ( ≥ 80% identity, ≥ 500 bp alignment) to avoid counting native background viral homologs as recovered spike-ins, and one phage was excluded from evaluation on the CST-IV-B background, where a native vaginal phage of the same host genus showed near-complete coverage at near-identical nucleotide identity to its spike-in counterpart.

**geNomad short-sequence reporting filter (ANI-novel panel)**

geNomad v1.11.2 reports a sequence shorter than 2,500 bp as viral only if it carries at least one virus hallmark gene (summary-module default `max_length_short_seq` = 2,500; run parameter `min_virus_hallmarks_short_seqs` = 1 and minimum score 0.7, both recorded in each run's summary JSON). The rule therefore applies to every 1,500-bp fragment and to no 3,000-bp fragment. To separate this reporting rule from classification, the aggregated virus score and virus-hallmark count of every ANI-novel fragment were read from geNomad's native aggregated-classification and marker-feature outputs and compared with its reported calls. At 1,500 bp, 45 of 65 homology-free and 331 of 427 homology-detectable fragments scored ≥ 0.7, but only the 4 and 134 carrying a hallmark were reported; every score-passing fragment that was not reported lacked a hallmark. At 3,000 bp, all score-passing fragments were reported (25 of 32 and 185 of 210), including 21 and 89 without a hallmark. No fragment below the cutoff was reported at either length. These counts are a diagnostic, not a replacement benchmark: bacterial negatives were not rescored, so they imply no relaxed-filter precision or MCC, and the 1,500- and 3,000-bp fragments are different sequences. The diagnostic script and per-fragment output are provided with the analysis code.

**Track B CST contrast: background provenance and scoring**

The primary 10× comparison used four CST-I and four CST-IV-B backgrounds. CST assignments were obtained from VIRGO2 profiles using VISTA and the VALENCIA traditional-CST classifier. CST-I included three runs from PRJNA1054643 (SRR27287961,

SRR27287984, SRR27287985) and one from PRJNA1356845 (SRR35936354). CST-IV-B included two runs from PRJNA1054643 (SRR27287964, SRR27287966), one from PRJNA1356845 (SRR35936363) and one from PRJNA1288683 (SRR34514383). Cohort representation was therefore unequal between the groups. Group means weighted backgrounds equally; 95% percentile intervals for ΔMCC (CST-IV-B minus CST-I) used independent resampling of backgrounds within each group (B = 2,000, random seed 12345; Table S9). Differences in precision and false-positive rate (FPR) used the same procedure, each metric with its own random-number generator (seed 12345), and FPR was computed as FP / (FP + TN) from the exact confusion counts.

The backgrounds differed in assembly size. After exclusions, the four CST-I backgrounds contributed 165 to 366 negative contigs and the four CST-IV-B backgrounds 9 to 2,975, so the viral fraction of scored contigs ranged from 24.7% to 41.9% in CST-I and from 3.7% to 92.5% in CST-IV-B. Because precision depends on this fraction, the precision difference was recomputed post hoc with each of the eight backgrounds omitted in turn (Table S9). For geNomad, VirSorter2 and ViraLM, omitting SRR27287964, the background with 2,975 negative contigs and a 3.7% viral fraction, reduced the difference to −0.013, −0.007 and −0.039, respectively.

Per-background exclusions removed contigs matching predefined native viral homologs of the spike-in panel before scoring. Spike-in-origin contigs were scored as positive and remaining background-origin contigs as negative. The exclusion procedure did not establish nonviral origin for all retained background contigs or comprehensively remove unrelated native viruses. Consequently, precision and MCC measure agreement with these operational labels; a call counted as false positive can include an unrecognised native virus. Because CST-IV assemblies in the independent cohort carried a larger labelled viral fraction (Table S39), unrecognised native viruses may be more frequent in CST-IV-B backgrounds, which would inflate their apparent false-positive counts.

For Track B, Sourmash calls required containment $\geq$ 0.5, and VirSorter categories 1–4 were counted as positive. DeepVirFinder and VirFinder calls required both score $\geq$ 0.5 and $p < 0.05$; AUPRC used the viral score alone. Scoring required successful tool execution and the expected result files. VirSorter identifiers were reconciled against assembly contig IDs, including circular-sequence annotations and prophage-region suffixes; multiple regions mapping to one parent contig contributed their highest category score. Unreported contigs were scored as negative, and per-category recall used the same binary calls as the overall metrics.

#### Kraken2 grounding of multi-evidence negatives

In the multi-evidence benchmark, Kraken2 assignments were parsed during ground-truth construction. Classified contigs were excluded from negative grounding if the reported taxon name contained "virus", "viridae" or "phage" (case-insensitive). Other classified names were retained, including unrecognised taxa. This operational parsing rule does not independently establish non-viral origin. Tier 0 additionally required no viral evidence line; CheckV host genes without viral genes provided the alternative grounding signal. Track C uses its separate bacterial/archaeal rule described above.

Because human assignments ground 2,833 of the 11,577 negatives (24%, and three times the 964 bacterial or archaeal assignments; Fig. S15B), we asked whether the tools misclassify them at a different rate. Each negative was assigned to one of the three grounding categories using the branch order above, and per-tool false positives were counted from the stored per-contig predictions at the documented operating thresholds (Table S21), so the counts are consistent with Table S12. False-positive rates on human-grounded contigs spanned 0% (VIBRANT and Sourmash) to 95.5% across the panel, and the ordering was not a rescaling of bacterial performance: TransGINmer misclassified 95.5% of human contigs as viral against 38.9% of bacterial or archaeal contigs (+56.6 percentage points), whereas Jaeger misclassified 0.9% against 20.1% (−19.2 points). Repeating the measurement on the 80 human-assigned Track C contigs in the background enrichment band, an independently labelled set, reproduced the ordering (Table S38). These are Kraken2 assignments rather than verified human sequence, contigs a tool did not report are scored negative, and Jaeger's 2,048 bp minimum input length restricts which negatives it reports; the comparison is therefore descriptive.

#### Question-specific stratifications (Q1–Q4)

Each benchmark question was tested with the metric appropriate to it. Q1 compared approach-level AUPRC at 500 bp and 1,000 bp. Q2 stratified Track A recall by virus category and tool scope. Q3 compared per-tool recall on the homology-free versus homology-detectable ANI-novel groups at each fragment length and detection of the Track C candidate dark-matter set. Q4 compared Tier-1 prophage recall and free-phage positive rates between a marker-dependent group (geNomad, VIBRANT, VirSorter, VirSorter2; n = 4) and a sequence-based group (DeepVirFinder, HVSeeker, MetaPhinder, PPR-Meta, TransGINmer, VirFinder, ViraLM; n = 7). All 49 Tier 1 prophage contigs carried homology (E1), CheckV (E2) and Phigaro (E3) support: 33 carried exactly these three lines, 10 also carried RCA support (E5), 3 CRISPR-spacer support (E4) and 3 both. A contig is categorised as a prophage only with Phigaro support, and Tier 1 requires at least three lines, so the set is restricted to prophages that homology searches, CheckV and

Phigaro already recognise, which may favour tools that use viral gene markers. The primary comparison uses the previously specified tool subset excluding Jaeger, Seeker and Sourmash, while values for all 14 tools are retained in Table S13. The comparison is conditional on that selection; Jaeger's minimum input length alone does not justify omitting it from a sufficiently long-contig analysis. Sourmash made no positive call on either contig class. Means were 91.3% versus 77.0% for prophage recall and 21.8% versus 37.0% for free-phage recall. Including Sourmash would lower the sequence-group means to 67.3% and 32.4%, respectively, without changing the direction of either difference. For each primary comparison, the Mann–Whitney U statistic was evaluated under all 330 allocations of four versus seven group labels, retaining tied ranks. Two-sided permutation p-values were twice the smaller inclusive tail probability, capped at one: $p = 0.097$ for prophages and $p = 0.073$ for free phages. Neither contrast was resolved at 0.05. These selected tools are not independent experimental interventions on marker dependence, and the comparison cannot isolate architecture, training and output filtering from the presence of marker features.

**Evidence-line dependencies and sensitivity checks**

E5 maps matched RCA reads to each shotgun contig and requires $\geq$ 10 read pairs at $\geq$ 95% identity over $\geq$ 70% of the contig. It measures read support, whereas Track C uses the library-size-normalised enrichment ratio on co-assembled contigs. MetaVR was constructed with geNomad v1.11.0, so E1b introduces an indirect dependency for geNomad. The three robustness analyses exclude E1b, merge E1 and E2, or exclude E5 (Table S11). When E1b is omitted, VIBRANT ranks first (MCC 0.448), ahead of geNomad (0.402); the full-evidence point estimates are nearly tied (0.439 and 0.440, respectively). This change shows that the ranking depends in part on the evidence definition and does not establish independence from shared-reference effects. Per-tier contributions are shown in Fig. S16. Across the $4 \times 4$ E5 threshold grid, TransGINmer ranks first in 15 of 16 settings and HVSeeker in the remaining setting (25% breadth and 41 read pairs). Top-five membership changes among six tools, including exchanges involving Seeker, PPR-Meta and VirFinder (Table S19; Fig. S9). This grid is separate from the Track C label sensitivity analyses.

A non-redundant E1b coverage estimator changes some E1b calls and tiers relative to the summed estimator used for the benchmark labels. Counts are given in the coverage-estimator sensitivity analysis below (Table S26).

**VMGC cross-check (annotation only)**

We compared the benchmark's novel sequences and reference databases against the vaginal-specific Vaginal Microbial Genome Collection (VMGC; Huang et al., 2024; Zenodo accession 10457006), using the full 14,224-genome set, the 4,263 vOTU representatives, and the `VMGC_virus.info` host and taxonomy table. VMGC uses the *Bifidobacterium vaginale* synonym for *Gardnerella vaginalis*; we report these jointly as *Gardnerella*-associated. Two comparisons were run. First, the 48 dark-matter contigs and the 15 ANI-novel panel genomes were searched against the full VMGC catalogue (BLASTn for all queries; skani for the full-genome panel). Second, the 4,263 vOTU representatives were searched against the two reference databases the benchmark uses for ground-truth annotation, MetaVR (BLASTn) and RefSeq viral proteins (DIAMOND BLASTx); these reference snapshots postdate the documented training corpora, so this comparison measures reference-database coverage rather than direct training-set overlap. A species-level (vOTU) match required $\geq$ 95% average nucleotide identity over $\geq$ 85% of the shorter sequence (Roux et al., 2019), computed with skani for full genomes and with a per-(query, subject) interval-merged BLASTn coverage estimator (below) for fragments and vOTUs; RefSeq protein homology required $\geq$ 30% identity over $\geq$ 50 amino acids at e $\leq$ 1e-10. VMGC was used purely as a post-hoc annotation layer and was never added to the multi-evidence ground-truth consensus, because VMGC was constructed with DeepVirFinder, VIBRANT, and CheckV, two evaluated tools and a tool used in ground-truth construction; adding it would reintroduce the same tool–ground-truth dependency that the E1b robustness checks already address. As an integrity check, the metadata join reproduced two independently published VMGC statistics exactly (2,814 of 4,263 vOTUs assigned to a viral family, 66%; 61 *Papillomaviridae* vOTUs; Huang et al., 2024). Of the 585 *Gardnerella*-associated VMGC vOTUs, 543 (92.8%) had no MetaVR species-level match. Host annotation does not establish whether related viral sequences are present: some sequence matches have unresolved or different predicted hosts. A direct BLASTn of the Track A *Gardnerella* phage (MW387018.1, *Gardnerella* phage vB_Gva_AB1) against MetaVR confirmed this present-but-unlabeled pattern at the sequence level: its best match was a species-level hit (97.0% ANI over 88% of the shorter genome; UViG IMGVR_UViG_3300008432_000002, with further hits at ~97% identity), yet that UViG is host-labelled only to family *Bifidobacteriaceae* with the genus unassigned, so the phage is present in MetaVR by sequence but is not annotated to its *Gardnerella* host. Per-vOTU and stratified overlap results are in Table S24, and the dark-matter and ANI-novel classifications in Table S25.

**MetaVR host-taxonomy query and its interpretation**

The host-annotation count used the MetaVR UViG metadata export (local copy dated 23 February 2026). The export contains 24,435,662 UViG records, of which 2,282,116 have a populated `host_taxonomy` field. Host lineages were parsed by rank. The selected rule retained exact species assignments to *Lactobacillus crispatus*, *L. gasseri*, *L. iners* or *L.*

*jensenii*, or genus assignments to *Atopobium*, *Mobiluncus*, *Fannyhessea* or *Sneathia*. These were 587 species matches plus 118 genus matches, totalling 705 records (0.0309% of populated host assignments). The recount script and full outputs are provided with the analysis code. This reconstructs a selected taxon rule; it does not establish that the rule was specified before inspecting the export.

Interpretation depends on the taxon set and annotation resolution. Using the whole *Lactobacillus* genus with the same four genera gives 3,089 records (0.1354%); additionally including *Prevotella*, *Megasphaera* and *Dialister* gives 53,549 (2.3465%). The broader genera include hosts from other body sites, so these are sensitivity analyses rather than estimates of vaginal origin. Among *Lactobacillus* assignments, 1,732 of 2,971 lacked a resolved species; 12,230 *Bifidobacterium* assignments also lacked species resolution. GTDB places the *Gardnerella* clade within *Bifidobacterium*. No record explicitly named a species of that clade, but the unresolved assignments preclude interpreting this as absence of *Gardnerella* phages. The VMGC and individual-phage sequence comparisons above demonstrate this distinction directly. The query counts host annotations in this export and cannot estimate the completeness of vaginal viral sequence coverage or a host-specific deficit in tool sensitivity.

**Mapping reported viral regions to evaluated contigs**

Each benchmark scores whether an evaluated parent contig contains a reported viral sequence. Direct contig identifiers were retained first. Tool-specific provirus, fragment or circular suffixes were removed only when the resulting parent ID matched an evaluated assembly contig; sample prefixes were retained so that identical assembly IDs from different samples could not merge. The maximum viral score, or the most confident reported viral category, was taken across the parent and all its reported regions. A reported geNomad provirus with virus score $\geq 0.7$ could therefore produce a positive parent call even when the surrounding whole-contig viral score was below 0.7. No additional whole-parent threshold was imposed on that tool. Nonviral annotations did not create positive calls. Parents without a qualifying direct or region call in a completed run were scored negative. VirSorter2 assigns no score to short sequences reported with the suffix `||lt2gene` (fewer than two genes); these were given a score of 0, consistent with their treatment as negative calls at the 0.5 threshold. Region detection is an operational contig-level outcome and does not assert that the entire parent sequence is viral. In fragment-results tables, an empty call set or a whole tool–length combination without score output has unavailable AUPRC; for binary all-tool comparisons, either is treated as making no positive calls. This convention includes Jaeger below its 2,048 bp minimum input length.

VIBRANT positives were taken from the final `.phages_combined.txt` identifiers, with the corresponding final combined FASTA as a fallback. Intermediate `VIBRANT_machine_*.tsv` labels were excluded because built-in curation can reject neural-network virus labels and accept sequences assigned other intermediate labels. An existing empty final call file represents zero accepted calls; intermediate files without a final call artifact are treated as incomplete output. The installed program reports version 1.0.1 despite its container tag of 0.5 (Table S21).

Where more than one completed VIBRANT run existed for an input (five UChoose samples and the Track C co-assembly), the run was selected from the completed-run logs, recorded input paths and output timestamps, without reference to benchmark labels. The input name and final phage count were checked against the log, and the selected final output is identified by its checksum in a manifest distributed with the analysis code.

**VIBRANT output stage in a published benchmark**

VIBRANT writes intermediate neural-network labels (`VIBRANT_machine_*.tsv`) before its curation step and reports its final calls in `.phages_combined.txt`. To test which output a published cross-biome benchmark scored, we reanalysed the gut data of Wu et al. (2024; Zenodo 10.5281/zenodo.10886947); the soil and seawater biomes were not examined. Their deposited per-contig calls reproduced the published gut VIBRANT metrics (their Table S8) for all eight virome–microbiome pairs. We reran VIBRANT v1.0.1 (container `multifractal/vibrant:0.5`, the image specified by the What_the_Phage v1.0.2 wrapper they used) with default settings in standard and virome mode on their contigs, after removing contigs present in both size fractions as they did (38,085 contigs), and kept both the intermediate and the final output. Among contigs labelled by both the deposited calls and our intermediate output (3,427 in standard and 3,509 in virome mode), no label conflicted, and our intermediate output reproduced the published true-positive rates to within 0.005. With the size fraction of origin as the label and metrics averaged over the eight pairs, the published standard-mode values (true-positive rate 0.177, false-positive rate 0.0150, precision 0.672) became 0.164, 0.0104 and 0.728 with the final calls, and the published virome-mode values (0.177, 0.0151, 0.672) became 0.255, 0.0474 and 0.512. Wu et al. report VIBRANT v1.2.1, whereas the specified container holds v1.0.1, so the version that produced their calls is uncertain; the label correspondence does not depend on it. About 8–10% of contigs were labelled by only one of the two runs, for reasons not resolved here. Size fraction is not a ground truth, because viromes carry microbial contamination and microbiomes carry prophages. We did not test whether these differences change the published tool ranking or apply to later VIBRANT releases. The rerun and comparison scripts are provided with the analysis code.

#### Sensitivity of evidence tiers to the Evidence-1b coverage estimator

The benchmark labels compute the Evidence-1b aligned fraction by summing BLASTn high-scoring-pair (HSP) alignment lengths per contig across all subject hits. This summed estimator can double-count overlapping query intervals and pool HSPs from different subjects, inflating the aligned fraction. To assess the effect on the labels, we also computed a non-redundant per-(query, subject) estimator that greedily tiles HSPs by descending bitscore, trims each to the query span not already covered by a higher-scoring HSP, and computes a length-weighted identity and aligned fraction over the non-redundant covered length; this estimator is also used for the VMGC cross-check. Applied to the same E1b BLASTn output for all 13 samples with the same thresholds ( ≥ 90% ANI, ≥ 75% aligned fraction), the non-redundant estimator would reclassify 4,307 contigs relative to the benchmark labels: 3,614 from Tier 3 (single-evidence, excluded from benchmarking) to Tier 0 (grounded negatives), 581 from Tier 3 to Tier −1 (neither viral nor non-viral evidence), and 111 from Tier 2 to Tier 3. Because the tier count combines protein and nucleotide homology into a single line (E1 = E1a OR E1b), a changed E1b call alters the tier only when no E1a hit is present; a single Tier-1 contig (a 1,398 bp contig lacking E1a support) would move to Tier 2. Recomputing the summed estimator from the same output reproduced the benchmark tier of every contig (100% concordance across all 13 samples). The benchmark uses the summed-estimator labels. VIBRANT ranked first when E1b was excluded, followed by geNomad (Table S11); that analysis does not determine the rankings under the non-redundant estimator. Per-sample counts are in Table S26.

#### External validation cohort: processing notes

The 30 external-validation metagenomes were selected from MiTCH (Shabana et al., 2026; ENA PRJEB108308), comprising a five-sample pilot and a 25-sample mgCST-stratified scale-up. Libraries were sequenced as 2 × 100 bp reads on DNBSEQ-T7 (MGI; Shabana et al., 2026). Reads were reprocessed with the primary-cohort benchmark pipeline, not the parent study's taxonomic-profiling workflow. The ground truth used four evidence lines (E1 homology, with E1a and E1b counted together; E2; E3; E4) and omitted the RCA cross-validation line (E5), which requires the paired virome data unavailable for these shotgun-only samples; rankings are therefore compared against the primary four-evidence (no-E5) baseline. The 14 tools were run once on the merged, sample-prefixed contig set after pre-selecting Tier 0, 1, and 2 contigs. Community state types were assigned de novo (VIRGO2 with the VISTA metagenomic-CST classifier, and VALENCIA for the traditional CST overlay), and the independently reproduced labels matched the stratified selection (25/25 scale-up samples). The diversity and per-community-state contrasts used a sample-block bootstrap (B = 2,000), with samples as the unit of replication.

**Sample and participant counts**

The primary UChoose analysis includes 13 paired shotgun–virome samples with 11 distinct participant-ID prefixes in `assets/samplesheet.csv`. UC055 contributes V1 and V2; UC093 contributes V2 and V3. Counts described as n = 13 therefore refer to samples, not independent participants. Contig-level uncertainty estimates do not account for all within-genome or repeated-participant dependence. The real shotgun-assembly analysis includes only contigs $\geq$ 1,500 bp; Fig. S1 shows that post-filter distribution and cannot establish the prevalence of shorter contigs in unfiltered assemblies.

Track C negatives require bacterial or archaeal classification and the background enrichment band; human assignments are excluded. This differs from the multi-evidence benchmark, whose negative grounding can include human assignments. Host depletion used hg19.

### Supplementary Figures

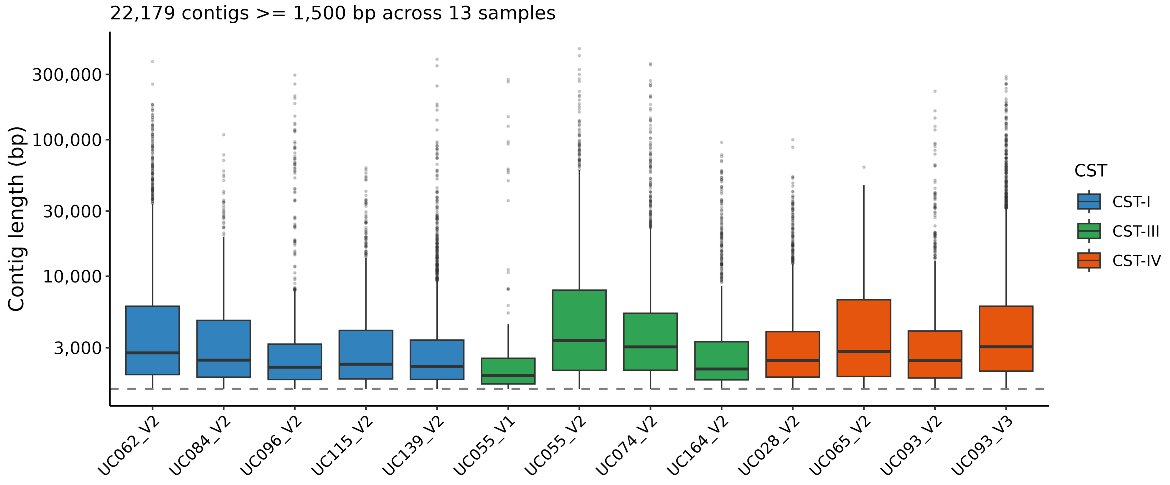


**Figure S1. Contig-length distribution of the primary UChoose shotgun-metagenome assemblies after the 1,500-bp filter.** Per-sample box plots of assembled contig length ($\log_{10}$ scale) for the 13 UChoose shotgun metagenomes that enter the multi-evidence benchmark, coloured by community state type, with the 1,500-bp inclusion threshold marked (dashed line). n = 22,179 contigs ≥ 1,500 bp across 13 samples.

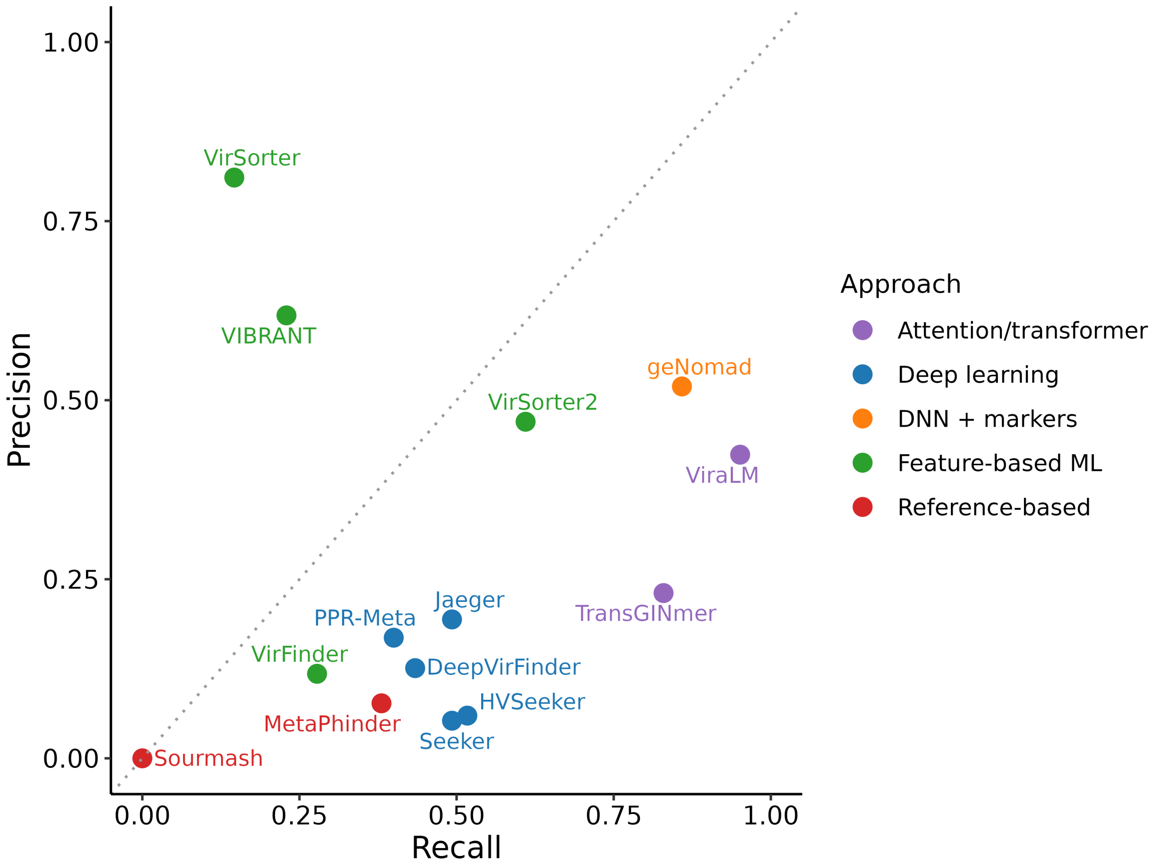


**Figure S2. Precision–recall scatter at 3,000 bp reproduces the 1,500-bp hierarchy with compressed spread.** Operating point of all 14 tools at 3,000 bp, each at its implemented operating threshold, in precision–recall space (Fig. 2B shows curves for the top six tools at 1,500 bp). n = 205 viral and 3,123 bacterial fragments.

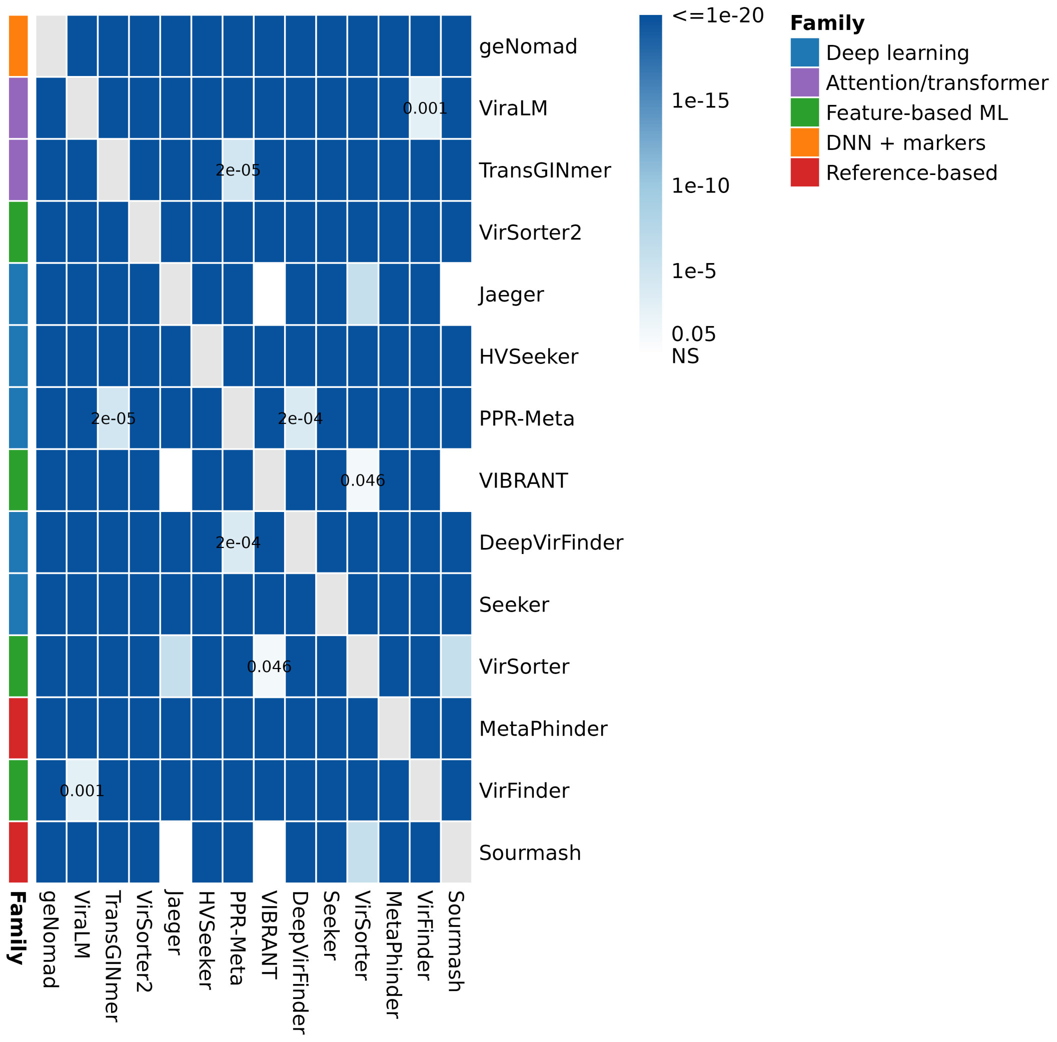


**Figure S3. Classification accuracy differs in 88 of 91 tool pairs at 1,500 bp.** Heatmap of pairwise McNemar q-values after Benjamini–Hochberg correction. Darker cells indicate stronger evidence of an accuracy difference; colour shows $-\log_{10} q$ capped at 20. White cells indicate non-significant comparisons ($q \geq 0.05$); the grey diagonal is untested. Sample size: 91 comparisons among 14 tools.

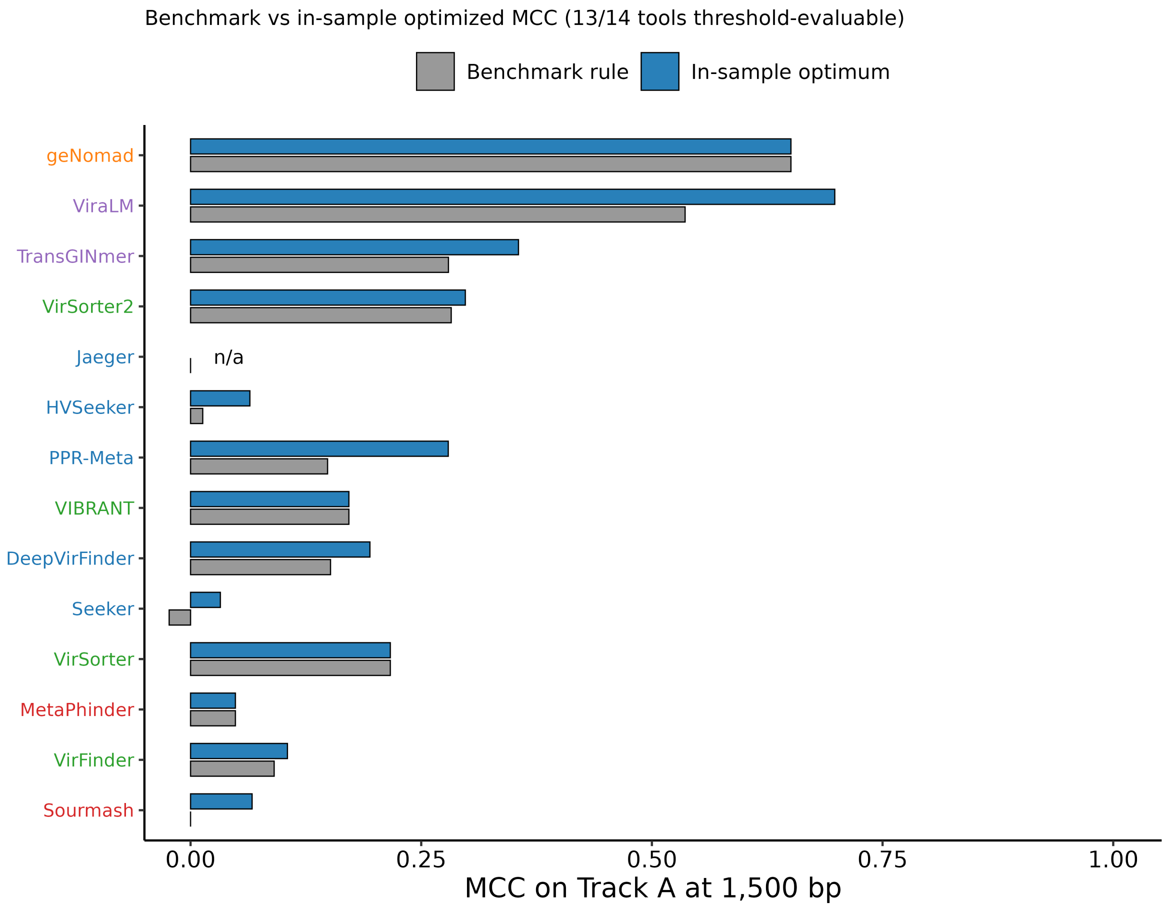


**Figure S4. In-sample threshold optimisation increases MCC for several tools at 1,500 bp.** Per-tool MCC under the implemented operating rule and the maximum obtained across candidate thresholds. Thresholds were selected and evaluated on the same fragments, so gains do not estimate out-of-sample improvement (Methods). Sample sizes: 6,689 fragments; 13 threshold-evaluable tools, with Jaeger unavailable at this length.

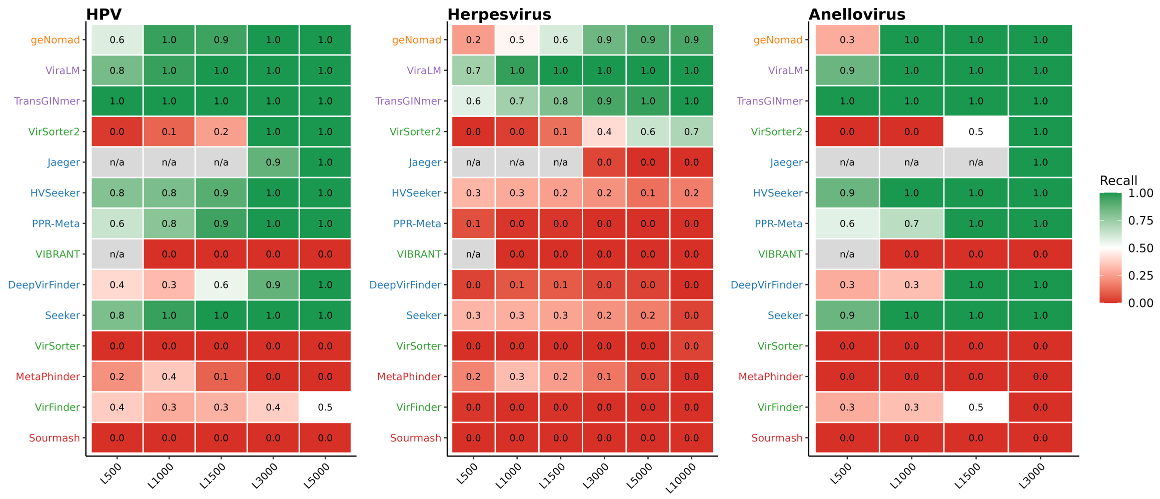


**Figure S5. Herpesvirus recall at 1,500 bp is complete only for ViraLM.** Per-tool recall stratified by eukaryote-infecting virus category (HPV, herpesvirus, anellovirus) across the six Track A fragment lengths. n = 4 HPV, 2 herpesvirus, and 1 anellovirus genomes.

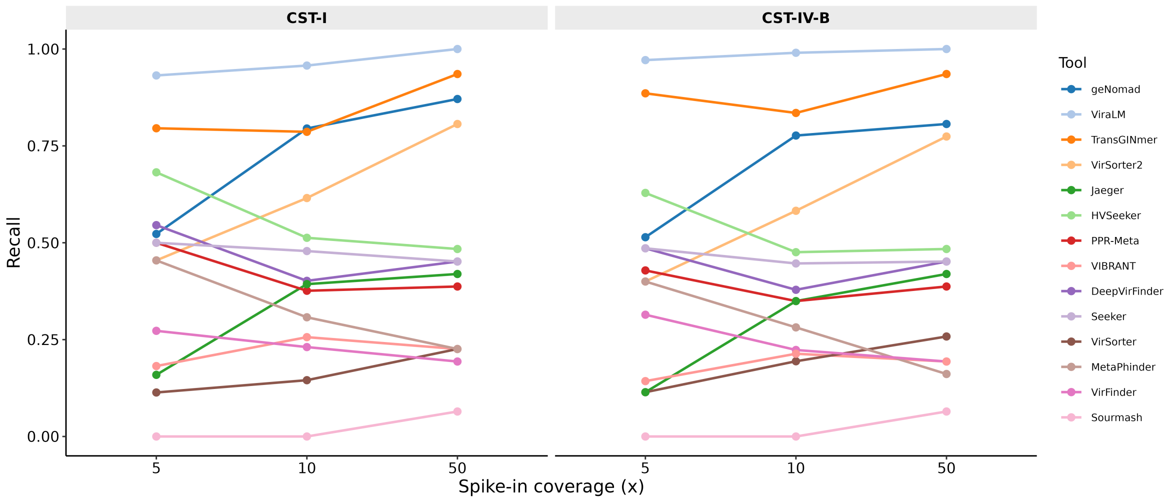


**Figure S6. Track B recall rises monotonically with simulated coverage for geNomad, ViraLM and**
**VirSorter2.** Per-tool recall on the two representative CST-I and CST-IV-B co-assemblies. No viral contig ≥ 1,500 bp assembles below 5×, so only 5×, 10× and 50× carry data. n = 14 tools × 3 coverage depths × 2 backgrounds.

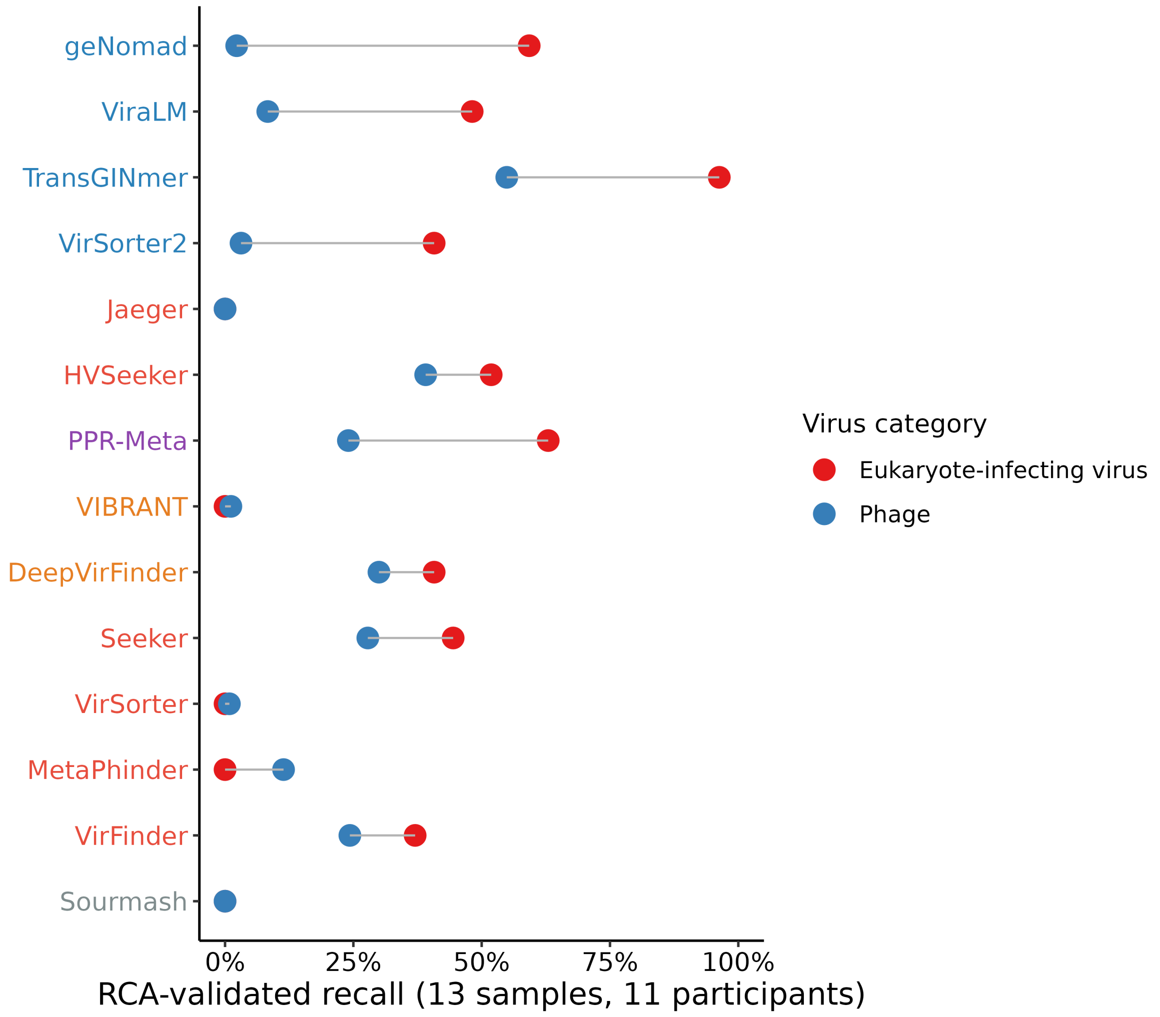


**Figure S7. RCA-validated recall is higher for eukaryote-infecting viruses than for phages in most tools.** Per-tool recall against RCA read support (E5), calculated from counts pooled across 13 shotgun assemblies and shown separately for phages and eukaryote-infecting viruses (Table S3; Supplementary Methods). Jaeger, Seeker and Sourmash returned no positive calls. Tool-label colours identify each tool's declared target scope (Table S21): phage-only (red), prokaryotic-virus (orange), all-virus (blue), phage+plasmid (purple) and database-dependent (grey). Sample sizes: 14 tools; 3,088 RCA-positive phage and 27 RCA-positive eukaryote-infecting virus contigs.

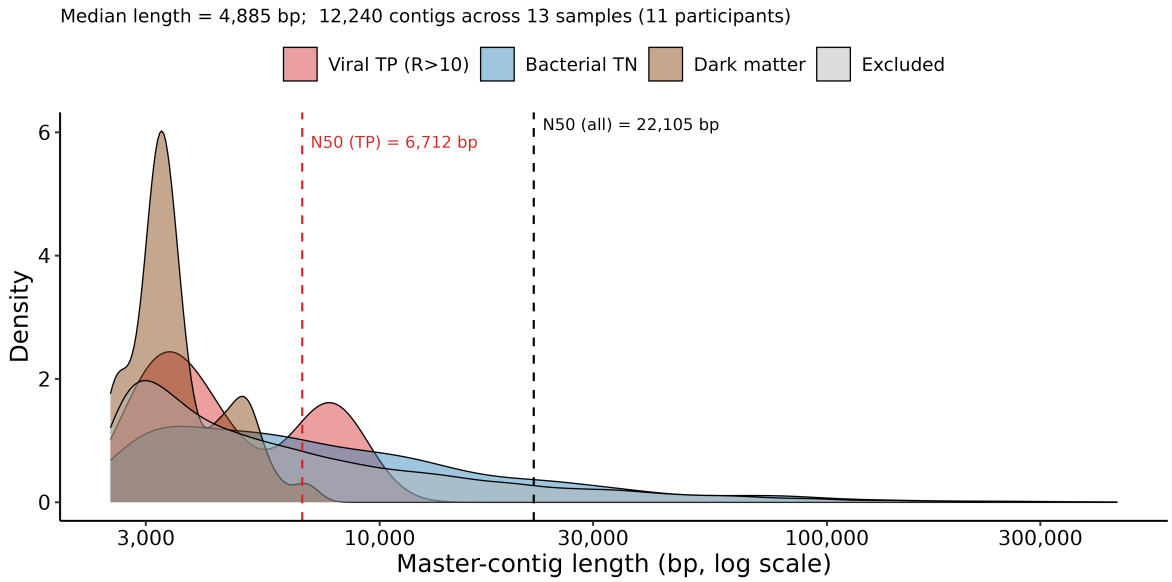


**Figure S8. Track C contig lengths vary by evaluation category.** Length distributions on a log scale for viral positives, bacterial negatives, candidate dark matter and excluded contigs. Categories combine enrichment and annotation evidence (Methods). Dashed lines mark N50 for all contigs (22,105 bp) and viral positives (6,712 bp). Sample sizes: 12,240 contigs ≥ 2,500 bp across 13 samples, including the 960-contig benchmark subset (29 positives and 931 negatives).

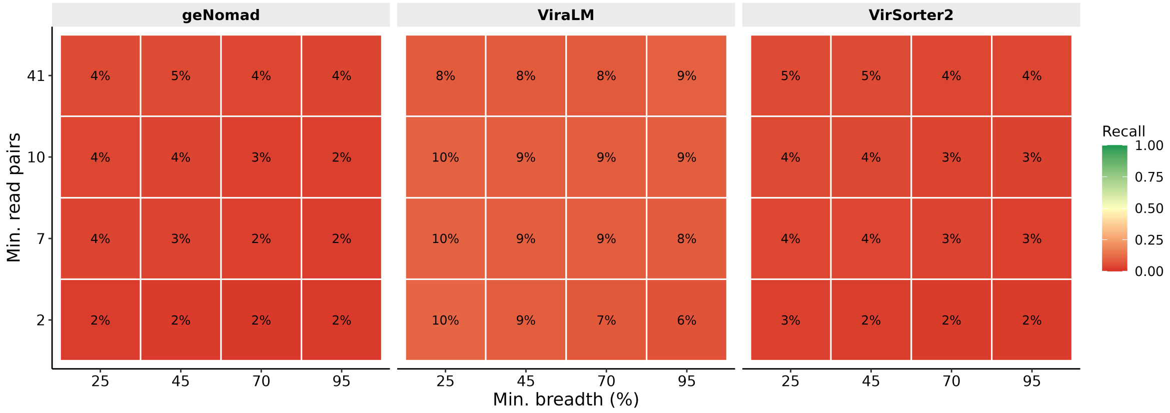


**Figure S9. RCA-supported recall depends on the Evidence-5 mapping thresholds.** Recall of geNomad, ViraLM, and VirSorter2 at 16 combinations of minimum mapping breadth and minimum paired-read count, computed over the shotgun contigs scored by Evidence line 5 of the multi-evidence benchmark. n = 3 tools × 16 threshold combinations; the RCA-positive contig set each cell defines ranges from 1,045 (95% breadth, ≥ 41 read pairs) to 14,004 (25% breadth, ≥ 2 read pairs).

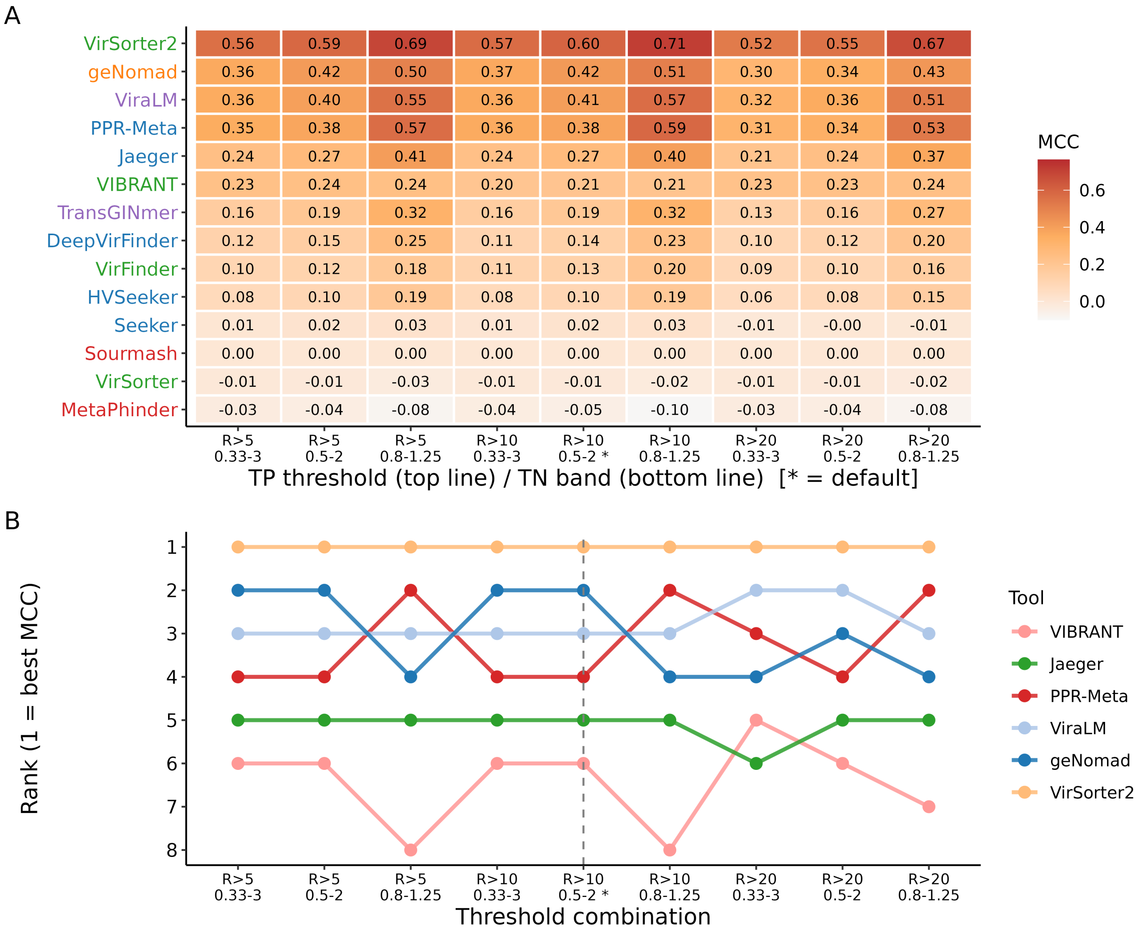


**Figure S10. VirSorter2 retains rank 1 on Track C across the 3 × 3 enrichment-ratio grid.** (A) Per-tool MCC at nine combinations of true-positive cutoff and true-negative band. (B) Per-cell ranking; VirSorter2 holds rank 1 in every cell and the same five tools compose the top five. n = 14 tools × 9 cells.

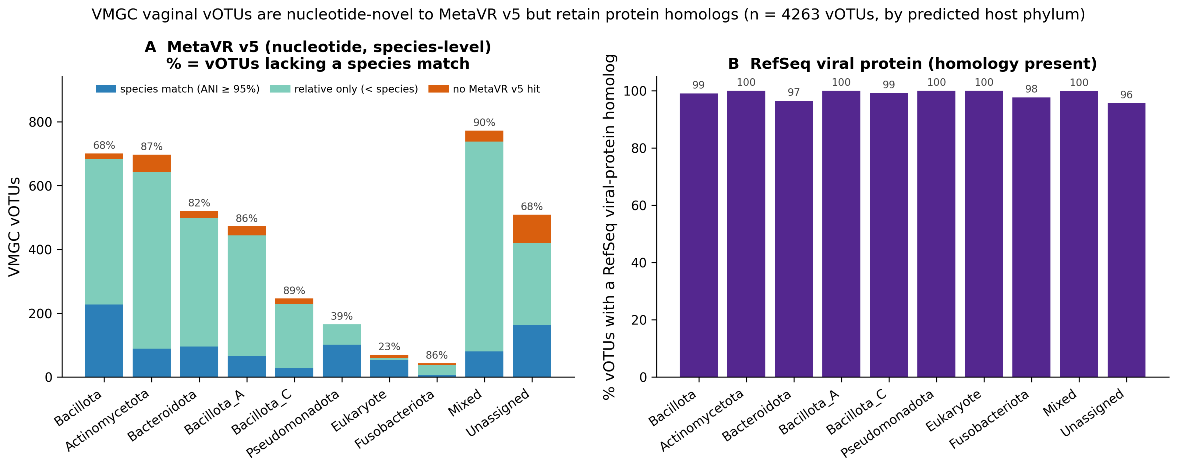


**Figure S11. Most vaginal vOTUs lack a species-level MetaVR match but retain protein homologs.** (A) MetaVR match categories by predicted host phylum: species-level match ( ≥ 95% ANI), more distant relative or no detectable hit. Labels give the percentage lacking a species-level match. Mixed denotes ambiguous multi-host predictions; Unassigned denotes no host assignment. (B) Percentage with detectable RefSeq viral-protein homology by host phylum. Search criteria are in Supplementary Methods. Sample size: 4,263 VMGC vOTUs.

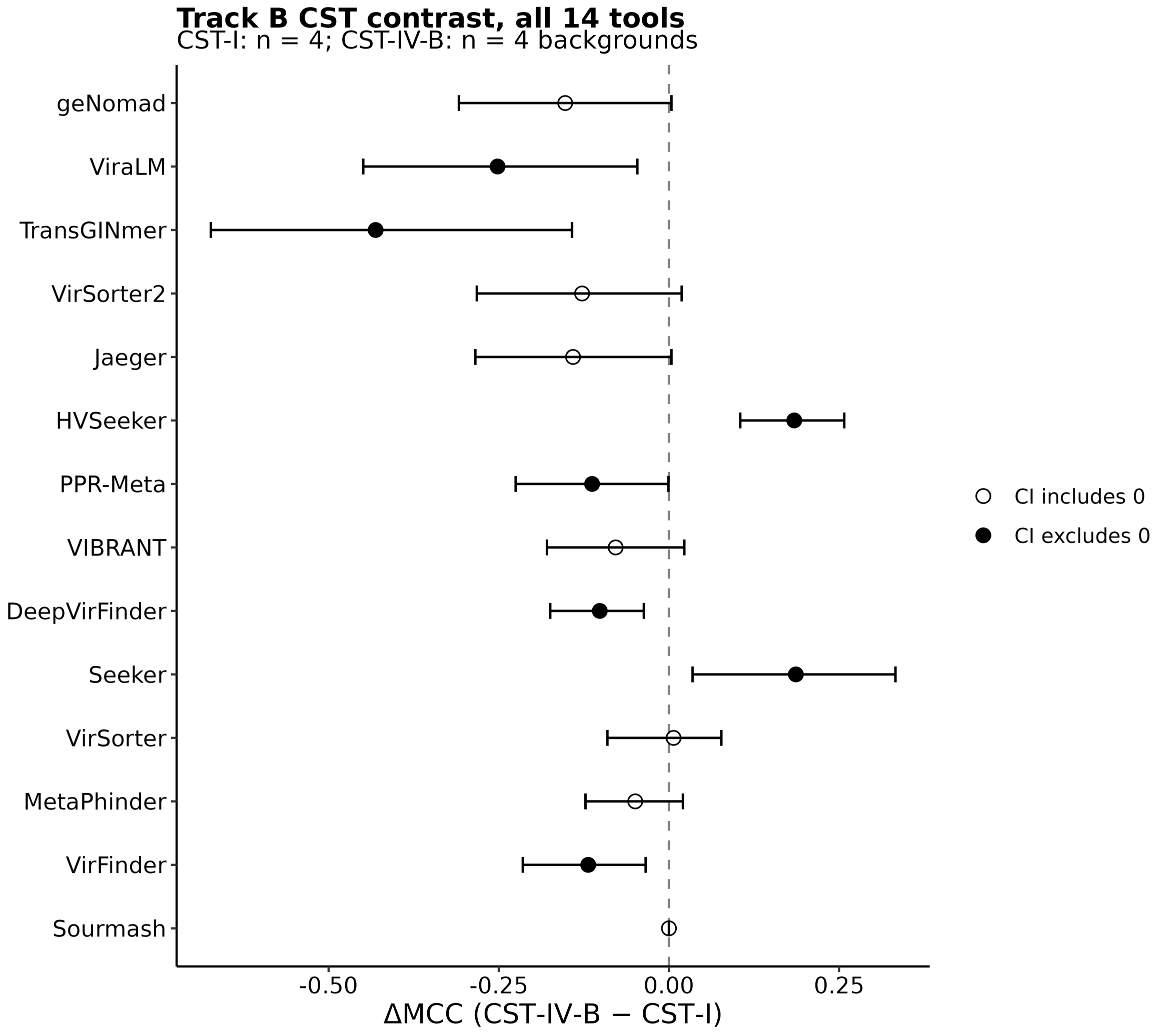


**Figure S12. CST-associated ΔMCC intervals support decreases for five tools and increases for two.** Per-tool ΔMCC (CST-IV-B minus CST-I) at 10× coverage, with background-level bootstrap 95% confidence intervals. Filled points indicate intervals excluding zero; open points indicate intervals spanning zero. Sample sizes: four backgrounds per CST group; 14 tools.

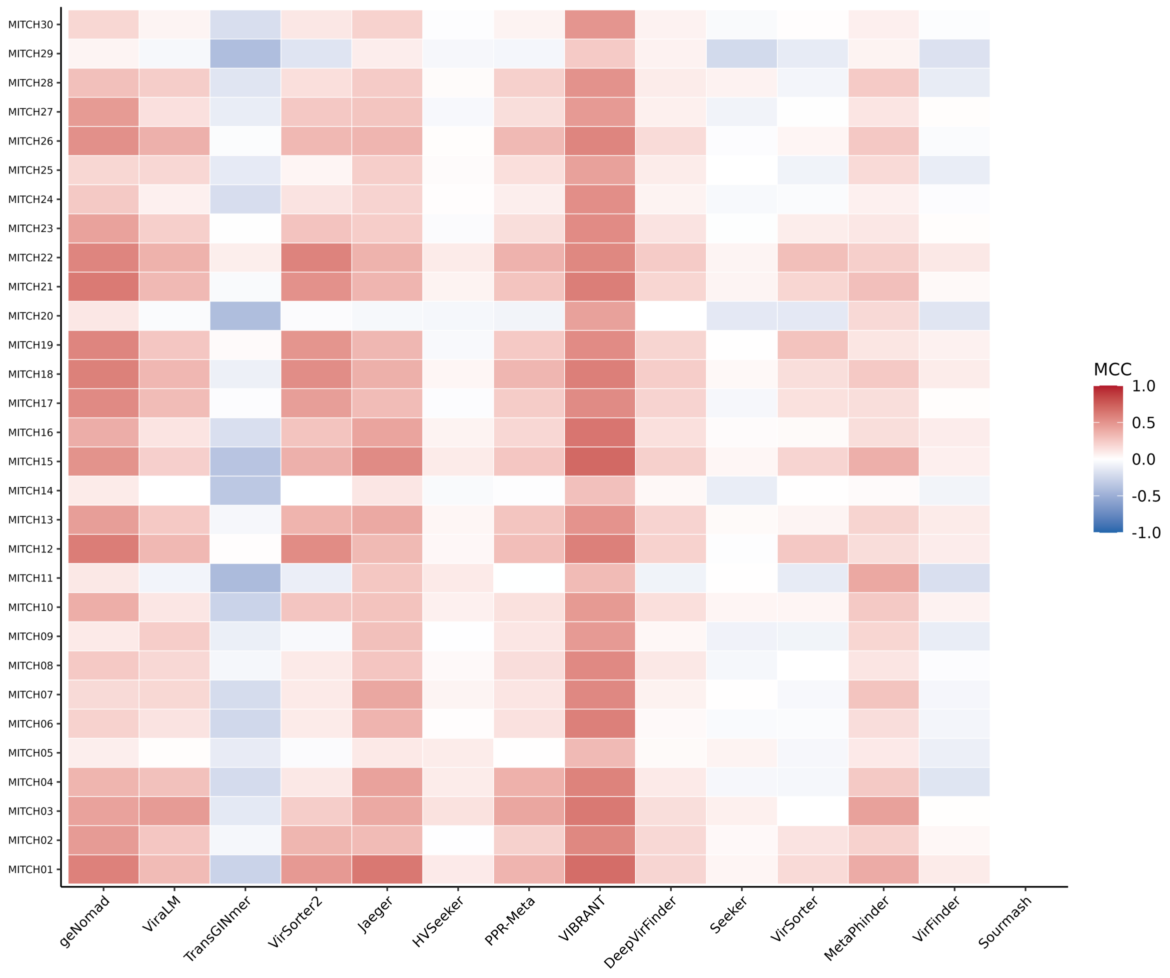


**Figure S13. Per-sample MCC matrix for the 30-sample external validation cohort.** Heatmap of per-tool MCC for each MiTCH shotgun metagenome (tools in columns, samples in rows), MCC on a blue-white-red scale centred at zero. n = 30 samples × 14 tools.

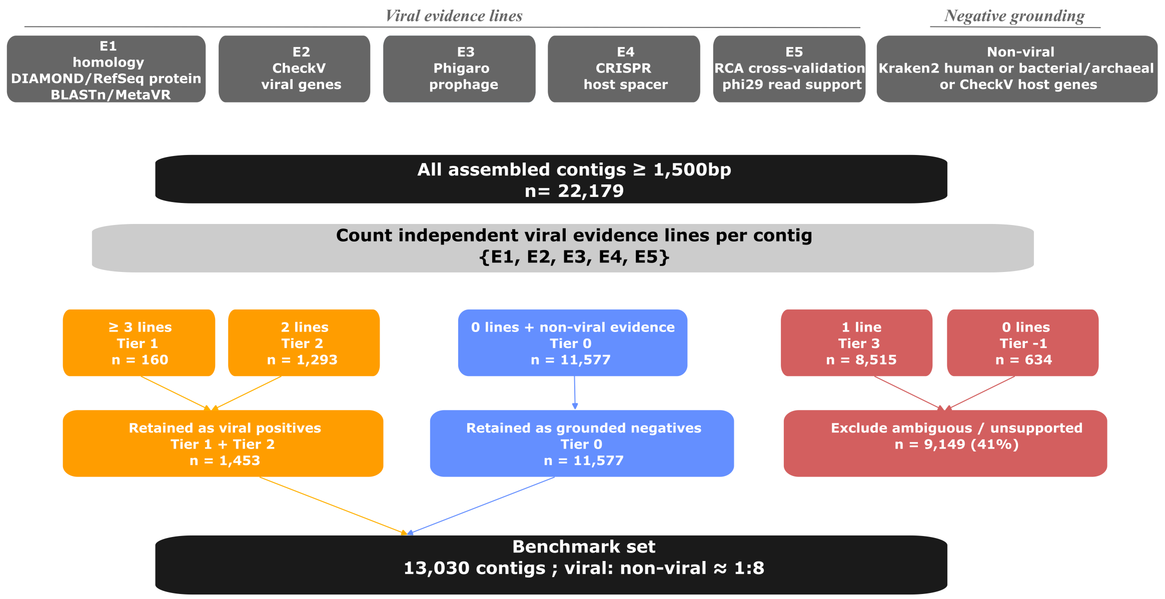


**Figure S14. Five evidence lines define the multi-evidence benchmark.** Contigs are assessed for homology (E1), structural signatures (E2), prophage integration (E3), CRISPR-spacer targeting (E4) and RCA read support (E5). Protein and nucleotide homology count as one line. Viral positives have at least three lines (Tier 1, n = 160) or exactly two (Tier 2, n = 1,293). Negatives lack viral evidence but have an explicit non-viral annotation (Tier 0, n = 11,577). Contigs with one viral evidence line (Tier 3, n = 8,515) or neither viral nor non-viral evidence (Tier −1, n = 634) are excluded. Thresholds and annotation rules are in Table S36 and Supplementary Methods. Sample sizes: 22,179 contigs ≥ 1,500 bp across 13 UChoose samples; 13,030 retained for evaluation.

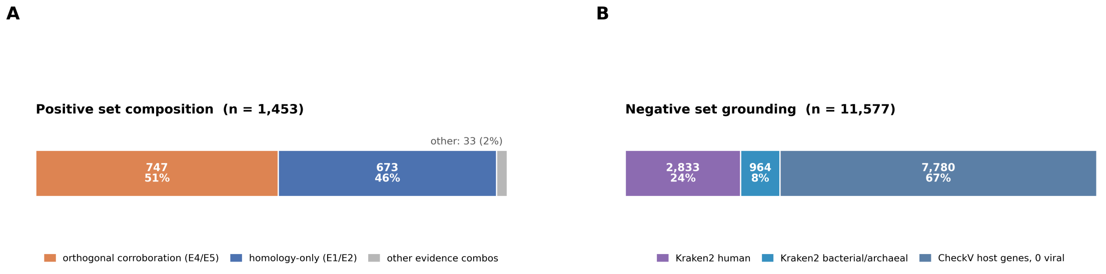


**Figure S15. Evidence supporting the retained positive and negative sets.** (A) Positive contigs supported by CRISPR or RCA evidence (747, 51%), E1/E2 reference-based evidence alone (673, 46%) or other combinations (33, 2%). (B) Negative contigs supported by CheckV host genes without viral genes (7,780, 67%), Kraken2 human assignments (2,833, 24%) or bacterial/archaeal assignments (964, 8%). Every negative requires explicit non-viral evidence (Fig. S14; Supplementary Methods). Sample sizes: 1,453 positives and 11,577 negatives from 13 UChoose assemblies.

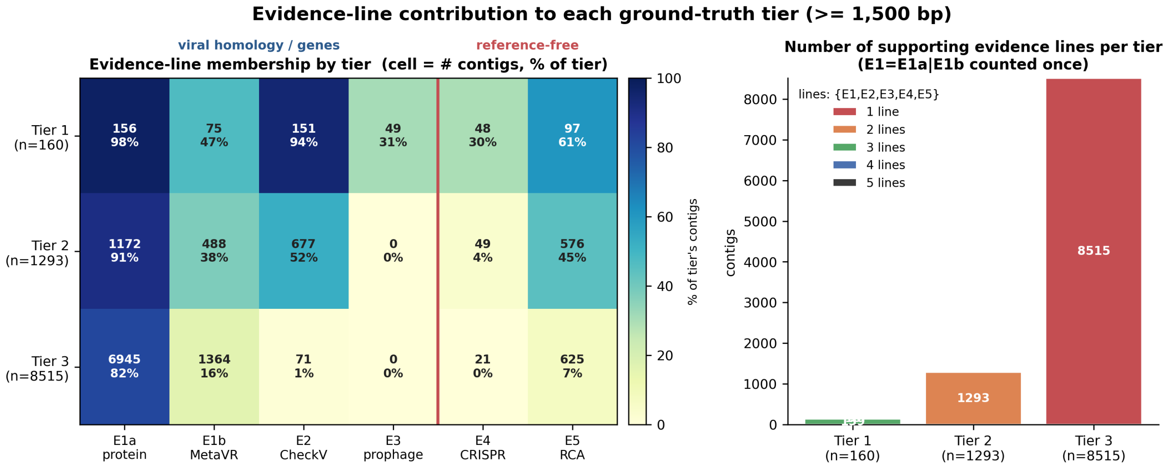


**Figure S16. Evidence-line contributions to each ground-truth tier.** (A) Heatmap of evidence-line membership by tier (Tier 1, n = 160; Tier 2, n = 1,293; Tier 3, n = 8,515; columns E1a protein, E1b nucleotide (MetaVR), E2 CheckV, E3 prophage, E4 CRISPR, E5 RCA), each cell the contig count and percentage of the tier. (B) Number of supporting evidence lines (E1a and E1b counted once) per tier; these lines are not assumed to be statistically independent. Based on the 13-sample pooled real-assembly contigs ≥ 1,500 bp.

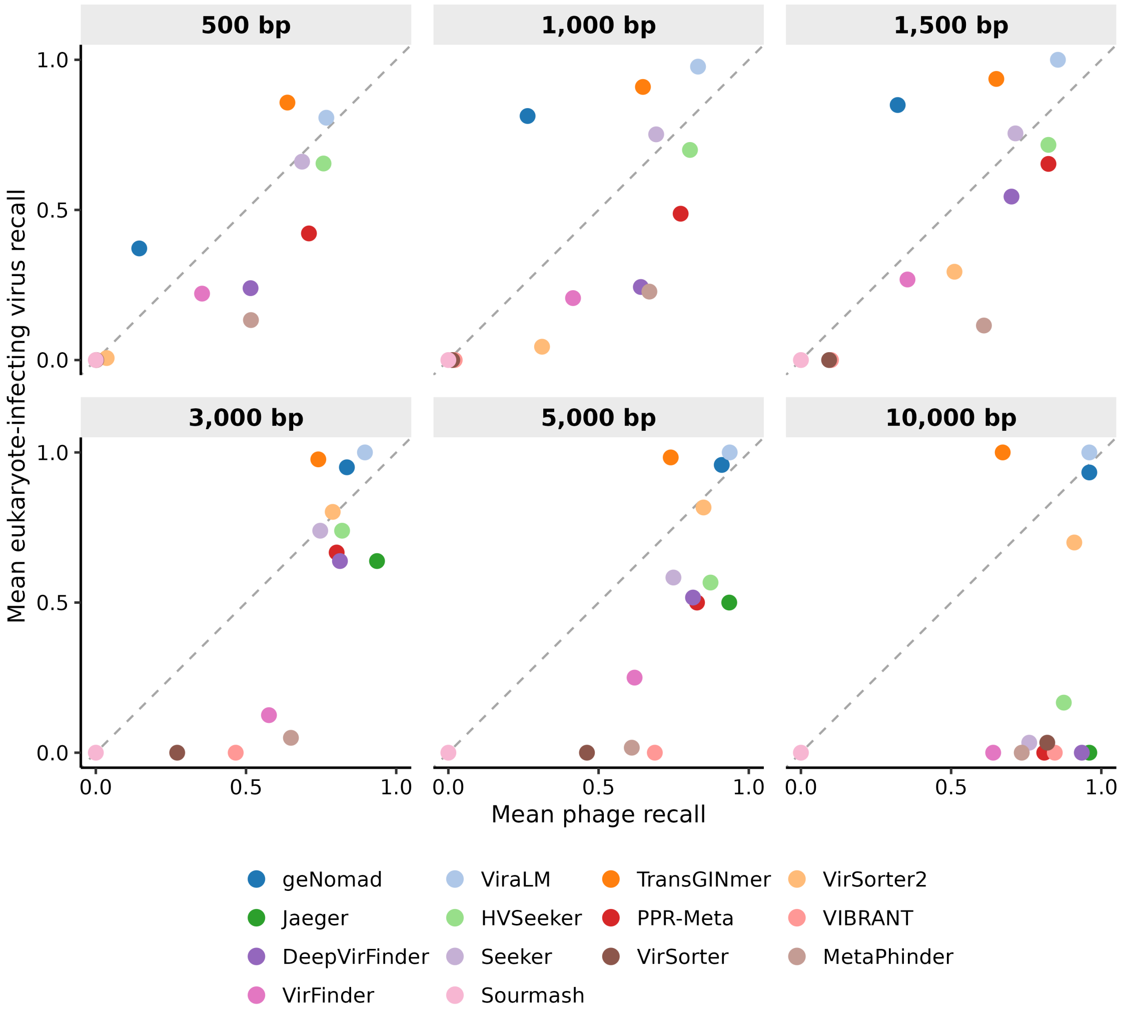


**Figure S17. Phage and eukaryote-infecting virus recall differ across tools and fragment lengths.** Each panel compares mean phage and eukaryote-infecting virus recall at one fragment length in the controlled-fragment benchmark (Track A). Colour identifies the tool; all panels share axes from 0 to 1, and the dashed diagonal marks equal mean recall. Group means give equal weight to each represented category: five phage host genera and up to three eukaryote-infecting virus categories. Anellovirus contributes no fragments at 5,000 or 10,000 bp, and human papillomavirus contributes none at 10,000 bp; the eukaryote-infecting virus mean at 10,000 bp therefore represents herpesviruses alone. Runs returning no classification are omitted (Jaeger at 500–1,500 bp and VIBRANT at 500 bp). n = 1,270, 630, 421, 205, 120 and 56 viral fragments at 500, 1,000, 1,500, 3,000, 5,000 and 10,000 bp, respectively; 12, 13, 13, 14, 14 and 14 evaluable tools at those lengths.

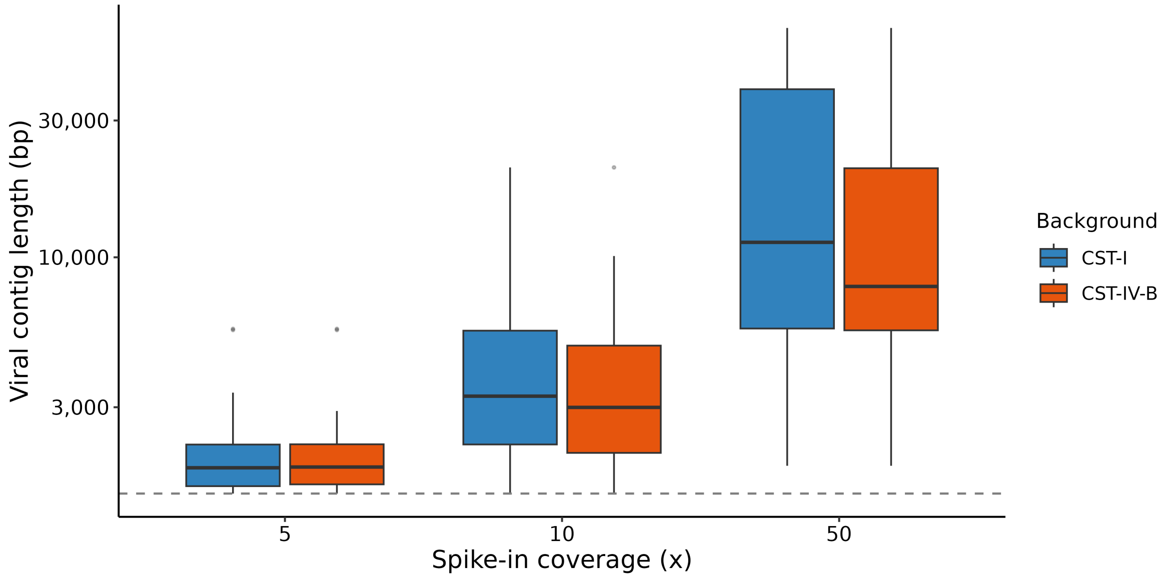


**Figure S18. Assembled viral-contig length increases with simulated coverage.** Contig-length distribution by per-genome coverage on the two representative CST-I and CST-IV-B backgrounds, with the 1,500-bp benchmark cutoff marked (dashed line). n = 361 viral contigs ≥ 500 bp across three coverage depths (5×, 10×, 50×) and two backgrounds.

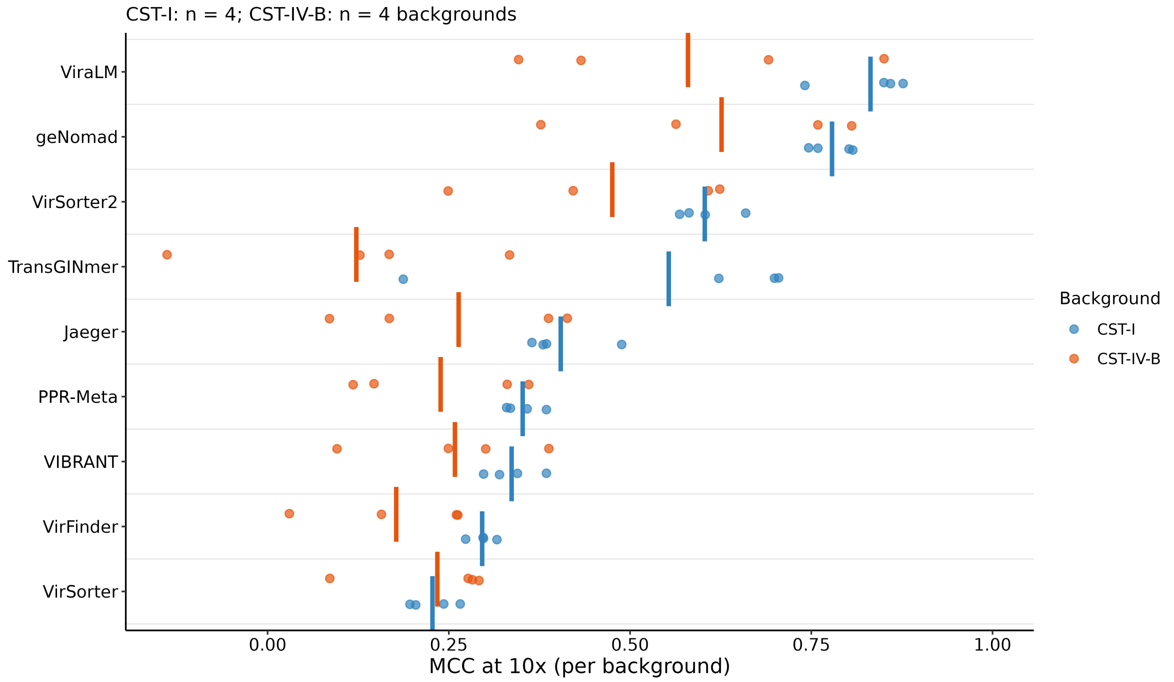


**Figure S19. Per-background MCC underlying the CST-IV-B diversity contrast.** MCC for the nine tools with the highest mean MCC in CST-I at 10× coverage across four CST-I and four CST-IV-B real backgrounds, with the CST groups distinguished by colour; each background is a point and the vertical bar is the arm mean. n = 4 backgrounds per arm.

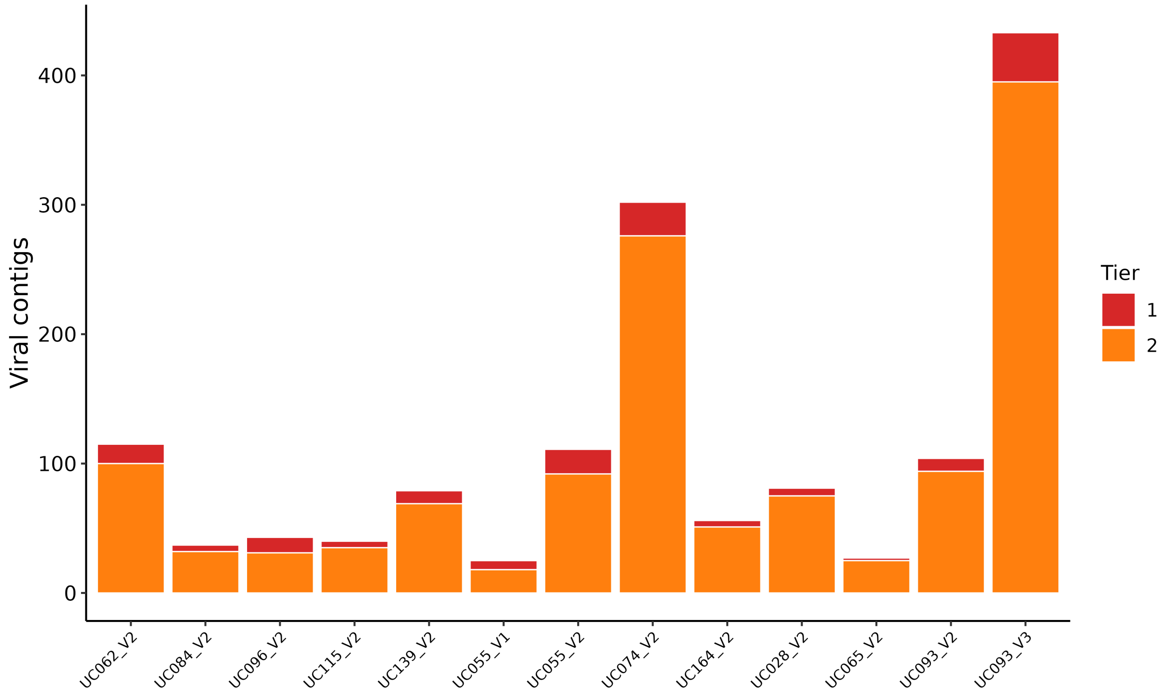


**Figure S20. Per-sample counts of viral-positive contigs by community state type.** Per-sample counts of Tier 1 and Tier 2 viral-positive contigs by CST across the 13 UChoose primary-cohort assemblies. n = 13 samples; 1,453 Tier 1+2 contigs.

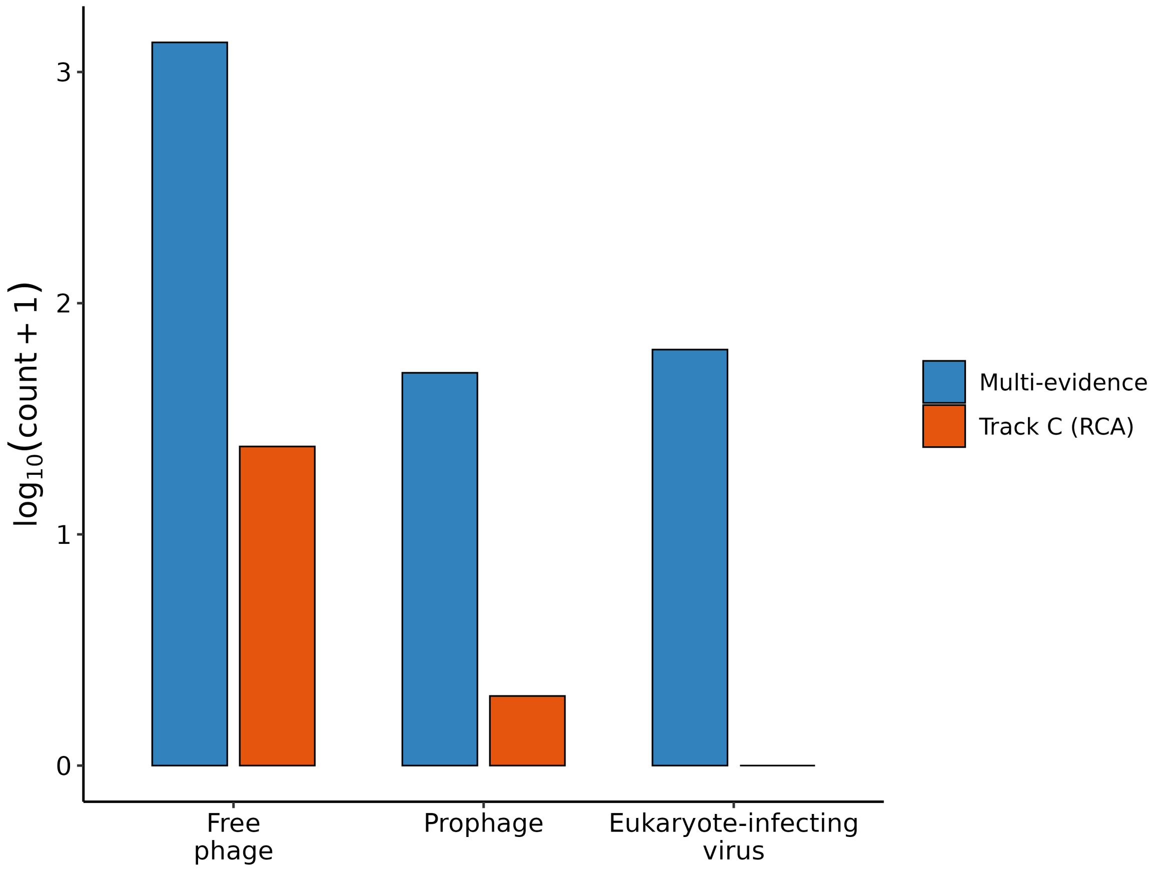


**Figure S21. Viral-category composition of the multi-evidence and Track C positive sets.** Viral-category composition of the multi-evidence Tier 1+2 viral-positive contigs and the Track C true-positive contigs (n = 29), on a $\log_{10}(\text{count} + 1)$ axis; free phage dominate both sets.

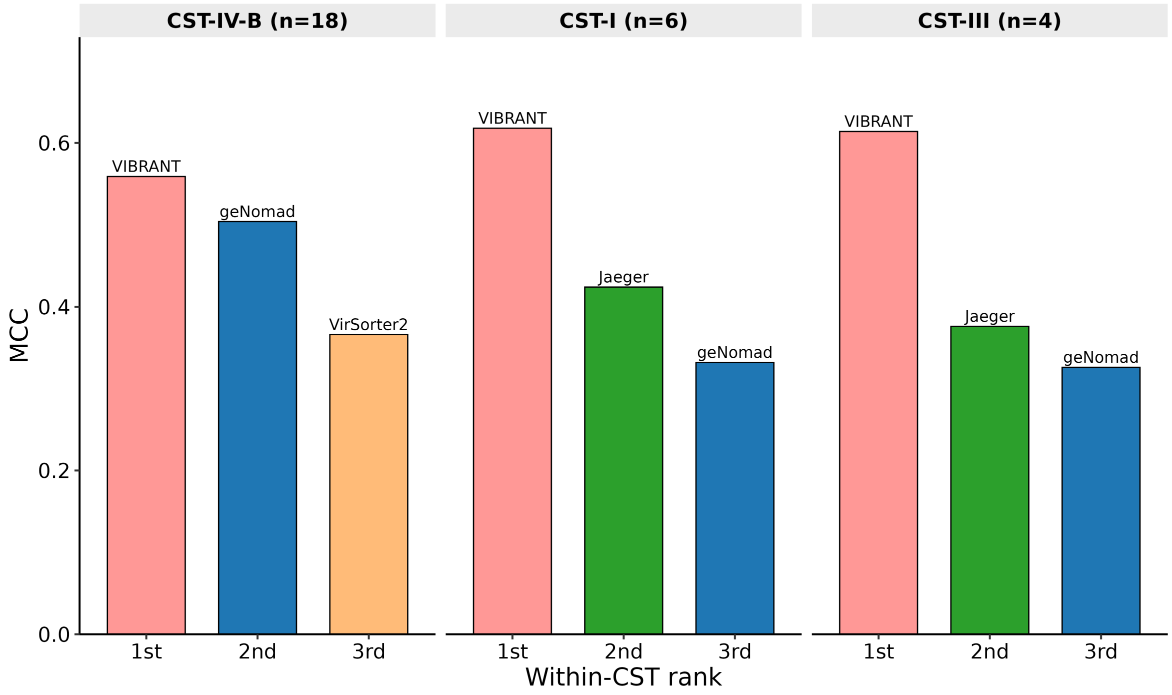


**Figure S22. Highest-MCC tools by community state type in MiTCH.** Results are shown for CST-IV-B (18 samples), CST-I (6) and CST-III (4). CST-V and CST-IV-C each contain one sample and are omitted. Comparisons within these small groups are descriptive. Sample size: 28 plotted samples from the 30-sample cohort.

### Supplementary Tables

Table data are in the accompanying workbook `supplementary_tables.xlsx`; each table below is one sheet, named by its table number.

**Table S1. Track A classification performance by virus category at 1,500 bp.** Per-category fragment counts, confusion-matrix counts and performance metrics for each tool, including false-positive rates on bacterial negatives. Sample sizes: 421 viral and 6,268 bacterial fragments; 14 tools.

**Table S2. Track C classification performance stratified by community state type, annotated with the constituent VISTA mgCSTs.** Per-tool confusion-matrix counts, recall, and precision on the Track C evaluation set within each community state type (CST-I, CST-III, CST-IV); the mgCSTs column lists the VISTA mgCSTs that map to each CST. n = 13 samples (5 CST-I, 4 CST-III, 4 CST-IV).

**Table S3. RCA-validated recall and precision by virus category across the 13 shotgun assemblies.** Per-sample and per-tool RCA-validated recall and precision with the underlying confusion-matrix cells, split into phage and eukaryote-infecting virus categories. n = 13 samples × 14 tools × 2 virus categories (364 rows).

**Table S4. Per-tool bootstrap 95% confidence intervals for every metric at every Track A fragment length.** 1,000-iteration bootstrap CIs for precision, recall, F1, MCC, and AUPRC. n = 84 tool × length cells.

**Table S5. Tool accuracy differs at every Track A fragment length.** Cochran's Q statistic, degrees of freedom and p-value for the comparison of per-fragment correctness across tools. Sample sizes: 14 tools at six fragment lengths.

**Table S6. Significant pairwise differences in Track A classification accuracy.** McNemar statistics and Benjamini–Hochberg-adjusted q-values for comparisons passing $q < 0.05$. Sample size: 485 significant comparisons among 546 tested (91 tool pairs at six fragment lengths).

**Table S7. Track A performance metrics by tool and fragment length.** Per-tool precision, recall, F1, MCC, and AUPRC at six lengths. n = 84 cells.

**Table S8. Track B performance across every coverage × representative-background combination.** Per-tool precision, recall, F1, MCC, AUPRC, and confusion-matrix counts. n = 14 tools × 3 coverages (5×, 10×, 50×) × 2 backgrounds (84 rows); the simulated depths below 5× yielded no viral contig ≥ 1,500 bp and are therefore absent from the table.

**Table S9. CST-I versus CST-IV-B performance at 10× coverage.** Per-group mean MCC, precision, recall and false-positive rate (FPR), with the CST-IV-B minus CST-I differences in MCC, precision and FPR and their background-level bootstrap 95% confidence intervals. Relative precision reduction is 100 × (CST-I – CST-IV-B) / CST-I. The post hoc leave-one-out columns give the smallest and largest precision difference when each of the eight backgrounds is omitted in turn, and the background whose omission gives the largest difference. Background-origin contigs were scored as negative after predefined native-homolog exclusions (Methods). Sample sizes: four backgrounds per group; 14 tools.

**Table S10. Track B per-tool recall by simulated coverage.** Recovery has a floor at 5× coverage. Per-tool recall restricted to viral contigs ≥ 1,500 bp in the two representative coverage-sweep backgrounds. n = 14 tools × 3 coverages (5×, 10×, 50×) × 2 backgrounds (84 rows); the simulated depths below 5× yielded no viral contig ≥ 1,500 bp and are therefore absent from the table.

**Table S11. MCC and rank under alternative evidence rules.** Per-tool results under the primary assignment, after excluding nucleotide homology (E1b), after merging homology and structural evidence (E1+E2), and after excluding RCA support (E5). Rules are detailed in Supplementary Methods. Sample sizes: 14 tools across four variants.

**Table S12. Per-tool confusion-matrix counts and metrics on the 13,030 pooled multi-evidence contigs.** Per-tool TP, FP, TN, FN, precision, recall, F1, MCC, and AUPRC on the benchmark-ready set ( ≥ 1,500 bp; Tier 0 negatives plus Tier 1–2 positives) pooled across all 13 samples, with 1,000-resample bootstrap 95% confidence intervals on MCC and AUPRC. n = 13,030 contigs (1,453 viral, 11,577 negative).

**Table S13. Prophage recall and free-phage positive rates by tool and marker dependence.** Per-tool results for all 14 tools, with the four marker-dependent and seven sequence-based tools included in the primary group comparison identified. Reported group means exclude Jaeger, Seeker and Sourmash; the Sourmash sensitivity analysis is described in Supplementary Methods. Two-sided permutation tests use the Mann–Whitney U statistic, retain tied ranks, and enumerate all 330 assignments of four versus seven tool labels; the tool is the unit of analysis. All 49 Tier 1 prophage contigs carried homology (E1), CheckV (E2) and Phigaro (E3) support, so the set is restricted to prophages these methods already recognise (Supplementary Methods). Sample sizes: 49 Tier 1 prophage and 1,342 free-phage contigs across 13 samples.

**Table S14. Per-contig evidence for the 48 dark-matter contigs.** Contig length, the library-size-normalised enrichment ratio $R$, and the number and identity of the tools detecting each contig. n = 48 contigs (median 3,317 bp, minimum 2,518 bp).

**Table S15. Per-tool detection of candidate dark-matter contigs.** Counts and fractions classified as viral under the implemented operating rules (Table S21). Sample sizes: 14 tools and 48 candidate contigs.

**Table S16. Runtime and peak memory on Track A and Track B.** Wall-clock runtime, peak RAM and GPU memory on the 1,500-bp Track A panel and the two representative 10× Track B co-assemblies (CST-I, 837 contigs; CST-IV-B, 5,149 contigs). Jaeger's 1,500-bp run returns no classification and is excluded from runtime comparisons. Values are single measurements under the documented hardware and settings (Methods). Sample sizes: 14 tools, one fragment panel and two co-assemblies.

**Table S17. geNomad virus scores on the 48 dark-matter contigs.** Lowering the virus-score threshold to 0.30 recovers only three of the 48. Per-contig geNomad virus score, plasmid score, and default-threshold classification. Median virus score across the set is 0.0001. n = 48 contigs.

**Table S18. Track C MCC and the derived per-tool ranking across a 3 × 3 grid of enrichment-ratio cutoffs.** Per-tool MCC at nine combinations of true-positive cutoff and true-negative band, with the per-cell ranking provided as a second sheet: VirSorter2 holds rank 1 in every cell and the same five tools (VirSorter2, geNomad, ViraLM, PPR-Meta, Jaeger) compose the top five, their order within ranks 2–5 varying. n = 14 tools × 9 cells.

**Table S19. RCA-validated recall and the derived recall-based ranking across the 4 × 4 Evidence-5 mapping-support grid.** Per-tool recall at 16 combinations of minimum mapping breadth (25%, 45%, 70%, 95%) and minimum paired-read count (2, 7, 10, 41), computed over the shotgun contigs scored by Evidence line 5 of the multi-evidence benchmark, with the recall-based per-cell ranking provided as a second sheet. First rank and top-five membership can change with the thresholds. n = 14 tools × 16 cells.

**Table S20. Per-source-sequence Track A recall and precision at 1,500 bp.** Per-sequence confusion-matrix counts, precision, and recall for the 14 viral source genomes and the 42 sequence accessions that make up the 6 bacterial negative-control type strains of Table S33 (one draft assembly contributes 36 of those contigs). n = 56 source sequences × 14 tools (784 rows).

**Table S21. Tool versions, operating rules and published scope.** Container details, training resources, methodological approaches and departures from default settings are provided for all 14 tools.

**Table S22. Tool rankings are broadly concordant across five evaluation contexts.** (A) Per-tool MCC in Track A at 1,500 and 3,000 bp, Track B at 10× in the CST-I coverage-sweep background, the multi-evidence benchmark, and Track C. (B) Pairwise Spearman correlations with raw and Benjamini–Hochberg-adjusted p-values. (C) The same comparisons restricted to the five highest-MCC tools in Track A at 1,500 bp (geNomad, ViraLM, VirSorter2, TransGINmer, VirSorter). (D) Sensitivity of the MiTCH rank-replication correlation to the tool set: Spearman rho against the matched no-E5 UChoose benchmark over all 14 tools, the eight tools with MiTCH MCC $\geq$ 0.10, and the five decision-relevant tools of Table 1, each with a sample-block bootstrap 95% confidence interval (B = 2,000), a permutation p-value (20,000 shuffles), and Lin's concordance correlation coefficient, which unlike rho penalises the systematic difference in MCC level between cohorts. Jaeger's unavailable 1,500-bp MCC is coded as zero in this correlation analysis. n = 14 tools in A/B, 5 in C; 10 pairwise comparisons.

**Table S23. Track C performance under alternative enrichment labels.** Per-tool MCC under labels derived from a three-component Gaussian mixture of $\log_{10}(R)$, compared with the primary enrichment cutoffs. Fitted component means, weights and Bayesian information criterion (BIC) are included; fitting and labelling rules are in Supplementary Methods. Sample sizes: 8,164 contigs with R > 0 and length $\geq$ 2,500 bp used for fitting; 14 tools evaluated.

**Table S24. MetaVR and RefSeq matches for VMGC vOTUs.** Per-vOTU nucleotide matches to MetaVR and protein homologs in RefSeq, annotated by viral family and predicted host. Summaries report match status by host phylum, host genus and viral family. Search thresholds are in Supplementary Methods. Sample size: 4,263 VMGC vOTU representatives.

**Table S25. VMGC matches for candidate dark matter and ANI-novel phage genomes.** Match status, matched vOTU, viral family and predicted host after searches against the full 14,224-genome VMGC catalogue. Contig matches require $\geq$ 95% identity over $\geq$ 50% of the contig; genome matches require $\geq$ 95% ANI over $\geq$ 85% of the shorter sequence. VMGC provides annotation only and does not determine benchmark labels (Supplementary Methods). Sample sizes: 48 candidate dark-matter contigs and 15 ANI-novel genomes.

**Table S26. Sensitivity of evidence tiers to the nucleotide-homology coverage estimator.** Per-sample E1b calls under the summed estimator used for the benchmark labels (columns marked orig or frozen) and under a non-redundant estimator (columns marked corrected), the resulting changes in evidence tiers, and reproduction of the benchmark tiers by the summed estimator. The non-redundant estimator removes overlapping alignments and separates reference hits (Supplementary Methods); the benchmark uses the summed-estimator labels. Sample size: 13 samples.

**Table S27. Per-tool pooled MCC in the external validation cohort and paired comparisons of the decision-relevant tools.** The ranking sheet reports pooled MCC with sample-block bootstrap 95% confidence intervals (B = 2,000), and the matched primary-cohort (four-evidence, no-E5) MCC and rank. The pairwise sheet reports all ten pairwise MCC differences among geNomad, VIBRANT, VirSorter2, ViraLM and Jaeger in the primary multi-evidence benchmark, Track C and MiTCH, with paired bootstrap 95% confidence intervals (B = 2,000). Contigs are resampled for the primary benchmark and Track C; samples are resampled for MiTCH. Intervals are nominal and unadjusted for multiple comparisons. Sample sizes: 14 tools across 30 MiTCH samples for rankings; 13,030 primary contigs, 960 Track C contigs and 30 MiTCH samples for paired contrasts.

**Table S28. Top-ranked tools within each traditional community state type in the external validation cohort.** Top three tools by MCC for each CST, with the number of samples per CST. The small CST groups are descriptive only. n = 30 samples (CST-IV-B = 18, CST-I = 6, CST-III = 4, CST-V = 1, CST-IV-C = 1).

**Table S29. Top-ranked tools within each metagenomic CST (mgCST) in the external validation cohort.** Top three tools by MCC for each mgCST, with the number of samples per mgCST. n = 30 samples.

**Table S30. Per-sample MCC matrix for the external validation cohort.** Per-tool MCC for each of the 30 MiTCH shotgun metagenomes. n = 30 samples × 14 tools.

**Table S31. High- versus low-diversity contrast in the external validation cohort.** Per-tool pooled MCC (confusion counts summed across samples before computing MCC) in high-diversity CST-IV samples (n = 19) and low-diversity CST-I/III/V samples (n = 11), the high-minus-low ΔMCC with sample-block bootstrap 95% confidence intervals (B = 2,000), and whether each interval excludes zero. n = 30 samples; 14 tools.

**Table S32. Per-sample community state type assignments for the external validation cohort.** De novo traditional CST (with subtype and confidence score) and metagenomic CST (mgCST, with score) for each of the 30 MiTCH samples, assigned with the VISTA classifier and Valencia (Methods). n = 30 samples.

**Table S33. The in silico spike-in reference panel.** Per-genome accession, organism, category, genome size, and host for the 14 viral and 6 bacterial genomes that define the Track A fragment pool. Sources: NCBI GenBank and MetaVR. n = 20 genomes.

**Table S34. Track A performance at 1,500 bp.** Per-tool precision, recall, F1, MCC, AUPRC and confusion-matrix counts, ordered by MCC. These are the 1,500-bp results from Table S7. Sample sizes: 14 tools and 6,689 fragments (421 viral, 6,268 bacterial).

**Table S35. Track C classification performance.** Per-tool confusion-matrix counts, precision, recall, false-discovery rate, F1, MCC and AUPRC, ordered by MCC, with bootstrap 95% confidence intervals for MCC and AUPRC. Methodological approaches and published target scopes (Table S21) are included. Sample sizes: 14 tools and 960 contigs (29 positives, 931 negatives).

**Table S36. Evidence sources and thresholds for the multi-evidence benchmark.** Biological signal, software, threshold and reference for protein and nucleotide homology (E1a/E1b), structural signatures (E2), prophage integration (E3), CRISPR-spacer targeting (E4), RCA read support (E5) and non-viral annotation. E1a and E1b count as one evidence line; tier rules are in Fig. S14. Sample size: 13,030 retained contigs ≥ 1,500 bp across 13 samples.

**Table S37. Recall by homology group and fragment length in the ANI-novel panel.** Per-tool true positives, false negatives, recall and recall differences between homology-detectable and homology-free genome groups at 1,500 and 3,000 bp (Fig. 5; Supplementary Methods). Tools are classified by dependence on external reference or marker databases; assignments are documented in sheet S37_taxonomy. Jaeger's 1,500-bp rows are flagged as unavailable and their zeros do not represent measured recall. The homology-detectable and homology-free groups contain different genomes, although all tools are evaluated on the same fragments within each group. Differences between genome groups therefore cannot be attributed solely to reference availability. Sample sizes: 14 tools × 2 groups × 2 lengths (56 rows); 13 homology-detectable genomes (427 and 210 fragments at 1,500 and 3,000 bp) and two homology-free genomes (65 and 32 fragments).

**Table S38. False-positive rate on the multi-evidence negatives differs by negative-grounding category.** Per-tool false positives and false-positive rates on the three categories that ground the multi-evidence negatives (Fig. S14; Fig. S15B): Kraken2 human assignments (n = 2,833), Kraken2 bacterial or archaeal assignments (n = 964), and CheckV host genes without viral genes (n = 7,780). Calls are the stored per-contig predictions evaluated at the documented operating thresholds (Table S21), so the counts are consistent with Table S12. The `contigs_reported_by_tool` column gives how many of the 11,577 negatives each tool returned a result for; contigs a tool did not report are scored negative under the benchmark rule, and Jaeger's lower count reflects its 2,048 bp minimum input length. The `human_minus_bact_arch_pp` column is the difference in percentage points, positive where a tool misclassifies human contigs more often. The final three columns repeat the measurement on the Track C human-assigned contigs falling in the background enrichment band, an independently labelled set (Methods). These are Kraken2 assignments, not verified human sequence, and the comparison is descriptive. Sample sizes: 11,577 negatives from 13 UChoose assemblies; 80 Track C contigs.

**Table S39. The CST-IV advantage in the external validation cohort depends on the MCC estimand and on viral prevalence.** Per-tool high-diversity CST-IV (n = 19) minus low-diversity CST-I/III/V (n = 11) ΔMCC under four estimands, each with a sample-block bootstrap 95% confidence interval (B = 2,000; the same draws as Table S31) and whether the interval excludes zero. `pooled` sums confusion counts across samples before computing MCC and reproduces Table S31. `mean_per_sample` averages per-sample MCC (Table S30), giving each sample equal weight. `prev_std_low` evaluates the CST-IV arm's pooled sensitivity and false-positive rate at the CST-I/III/V labelled viral fraction; `prev_std_cohort` evaluates both arms at the whole-cohort fraction. The prevalence-standardised estimands hold each arm's sensitivity and false-positive rate fixed and adjust the metric only, not the biology. Labelled viral fractions are 2.0% (CST-I/III/V), 6.4% (CST-IV) and 4.7% (cohort); they describe the evaluated contig sets ( ≥ 1,500 bp, four-evidence labels), not measured viral abundance. Intervals are nominal and unadjusted for the 14 tools and four estimands. n = 30 samples; 14 tools.

**Table S40. Bacterial diversity, labelled viral contigs and per-sample MCC in the external validation cohort.** (A) Per sample: traditional CST and contrast arm, bacterial Shannon diversity, richness (taxa at ≥ 0.1% relative abundance) and *Lactobacillus* fraction from VIRGO2 taxon abundances; evaluated contigs, labelled viral contigs by category (free phage, prophage, eukaryote-infecting virus) and labelled viral fraction; and MCC for each tool (as Table S30). (B) Spearman correlations between bacterial Shannon diversity and labelled viral contig count, viral fraction and contig count, and between each tool's per-sample MCC and viral fraction or Shannon diversity, for all samples and for CST-IV-B samples only, with bootstrap 95% confidence intervals (B = 2,000) and two-sided permutation p-values (10,000 permutations). Hosts were not assigned to contigs, so the correlations do not identify why viral contigs are more numerous in diverse communities. Analyses are exploratory and unadjusted for multiple testing. n = 30 samples (18 CST-IV-B); 14 tools.

**Table S41. Threshold optimisation estimates depend on the evaluation fragments.** Per-tool MCC at the documented benchmark operating rule and at the score cutoff that maximises MCC on the same Track A fragments, with both cutoffs and their difference. Results are reported at 1,500 and 3,000 bp; missing values identify runs without threshold-evaluable scores. Binary final calls are evaluated on their available score levels, and filtered outputs do not represent unrestricted classifier scores. These optima are descriptive in-sample estimates and do not establish transferable operating settings. Figure S4 displays the 1,500-bp comparison. Sample sizes: 14 tools at two lengths; 6,689 fragments at 1,500 bp and 3,328 at 3,000 bp.
